# PCH-2^TRIP13^ regulates spindle checkpoint strength

**DOI:** 10.1101/389080

**Authors:** Lénaïg Défachelles, Anna E. Russo, Christian R. Nelson, Needhi Bhalla

## Abstract

Spindle checkpoint strength is dictated by three criteria: the number of unattached kinetochores, cell volume and cell fate. We show that the conserved AAA-ATPase, PCH-2/TRIP13, which remodels the checkpoint effector Mad2 from an active conformation to an inactive one, controls checkpoint strength in *C. elegans*. When we manipulate embryos to decrease cell volume, PCH-2 is no longer required for the spindle checkpoint or recruitment of Mad2 at unattached kinetochores. This role in checkpoint strength is not limited to large cells: the stronger checkpoint in germline precursor cells also depends on PCH-2. PCH-2 is enriched in germline precursor cells and this enrichment relies on conserved factors that induce asymmetry in the early embryo. Finally, the stronger checkpoint in germline precursor cells is regulated by CMT-1, the ortholog of p31^comet^, which is required for both PCH-2’s localization to unattached kinetochores and its enrichment in germline precursor cells. Thus, PCH-2, likely by regulating the availability of inactive Mad2 at and near unattached kinetochores, governs checkpoint strength. This role may be specifically relevant in scenarios where maintaining genomic stability is particularly challenging, such as in oocytes and early embryos enlarged for developmental competence and germline cells that maintain immortality.

## INTRODUCTION

To prevent the missegregation of chromosomes and the production of daughter cells with an incorrect number of chromosomes, the spindle checkpoint (also called the spindle assembly checkpoint or the mitotic checkpoint) monitors whether chromosomes are attached to the spindle via kinetochores. If kinetochores fail to attach properly, this checkpoint delays the cell cycle to promote error correction and prevent aneuploidy. Despite its critical role, the duration of the cell cycle delay, defined as the strength of the spindle checkpoint, can be highly variable. This variability can be controlled by the number of unattached kinetochores (Collin *et al*., 2013), cell volume (Galli and Morgan, 2016; Kyogoku and Kitajima, 2017), and cell fate (Galli and Morgan, 2016; Gerhold *et al*., 2018).

The spindle checkpoint response initiates with the recruitment of Mad1 and Mad2 at unattached kinetochores (Chen *et al*., 1996; Li and Benezra, 1996; Chen *et al*., 1998; Sironi *et al*., 2001), which catalyzes the production of a Mitotic Checkpoint Complex (MCC). The MCC enforces a checkpoint arrest by inhibiting the Anaphase Promoting Complex/Cyclosome (APC/C) and preventing cell cycle progression (Sudakin *et al*., 2001). Formation of the MCC is driven by conformational changes in Mad2, which can exist in an open conformation (O-Mad2) or a closed conformation (C-Mad2) (Luo *et al*., 2002; Sironi *et al*., 2002; Luo *et al*., 2004). Mad2 is in the closed conformation in the Mad1/Mad2 tetramer recruited to unattached kinetochores. C-Mad2 in the tetramer acts as a template to convert additional soluble O-Mad2 to C-Mad2, which can be assembled into the MCC (Sironi *et al*., 2001; De Antoni *et al*., 2005; Simonetta *et al*., 2009; Fava *et al*., 2011). Thus, unattached kinetochores act as a platform for MCC assembly. This soluble signal generated by unattached kinetochores effectively tunes the spindle checkpoint response: the length of the cell cycle delay imposed by the checkpoint is governed by the ratio of unattached kinetochores producing MCC, and its ability to inhibit the APC, to cytoplasmic volume (Collin *et al*., 2013; Dick and Gerlich, 2013; Galli and Morgan, 2016; Kyogoku and Kitajima, 2017).

PCH-2/TRIP13 is a hexameric AAA+ ATPase that remodels HORMA domain-containing proteins, a group that includes Mad2 (Aravind and Koonin, 1998; Rosenberg and Corbett, 2015; Vader, 2015). Biochemical and structural studies have shown that PCH-2 converts C-Mad2 to O-Mad2 (Ye *et al*., 2015; Brulotte *et al*., 2017; Alfieri *et al*., 2018). TRIP13 works with the adaptor protein p31^comet^ to extract C-Mad2 from the MCC and promote its disassembly, permitting the activation of the APC/C and silencing the checkpoint (Eytan *et al*., 2014; Wang *et al*., 2014; Miniowitz-Shemtov *et al*., 2015; Ye *et al*., 2015; Brulotte *et al*., 2017; Alfieri *et al*., 2018). In addition to this role, we and others have shown that PCH-2/TRIP13 is essential for spindle checkpoint activation in *C. elegans* and mammalian cells (Nelson *et al*., 2015; Ma and Poon, 2016; Yost *et al*., 2017; Ma and Poon, 2018). PCH-2 is present at unattached kinetochores (Tipton *et al*., 2012; Wang *et al*., 2014; Nelson *et al*., 2015) and is required to robustly localize Mad2, but not Mad1, to unattached kinetochores (Nelson *et al*., 2015; Yost *et al*., 2017). A major implication of this work is that O-Mad2 can be limiting during checkpoint activation and PCH-2/TRIP13 plays a central role in ensuring its availability (Ma and Poon, 2018).

Based on a genetic interaction between the *C. elegans* ortholog of p31^comet^, CMT-1, and PCH-2, we had previously proposed that PCH-2 disassembles a CMT-1/Mad2 complex to promote checkpoint signaling, similar to its role during checkpoint silencing (Nelson *et al*., 2015). However, recent data from mammalian systems, in which loss of p31^comet^ does not suppress the requirement for TRIP13 (Nelson *et al*., 2015; Ma and Poon, 2016; Yost *et al*., 2017; Ma and Poon, 2018) and TRIP13’s function becomes essential for checkpoint activity only when O-Mad2 becomes limiting (Ma and Poon, 2018), suggest an alternative model in *C. elegans*. In this model, some Mad2, likely C-Mad2, is complexed with and stabilized by CMT-1, explaining the reduction in total Mad2 protein levels in *cmt-1* mutants. Despite this reduction of Mad2, we propose that in the absence of CMT-1, more O-Mad2 is available, thus making PCH-2 partially dispensable and explaining the genetic suppression. This model differs from our understanding of TRIP13 and p31^comet^ in mammalian cells, potentially because relative levels of C and O-Mad2 may vary between systems and most Mad2 in mammalian cells is present as O-Mad2 (Luo *et al*., 2004). Further, it highlights the importance of studying spindle checkpoint function in developmentally-relevant model organisms.

This model, however, raises another question: if the primary role of PCH-2/TRIP13 is to guarantee enough O-Mad2 is available for checkpoint activation and this role can be dispensable when enough O-Mad2 is available, is there a reason for PCH-2/TRIP13 to localize to unattached kinetochores (Tipton *et al*., 2012; Wang *et al*., 2014; Nelson *et al*., 2015)? One possible answer comes from our additional analysis of *cmt-1* mutant worms. In addition to its role as a PCH-2 adapter (Ye *et al*., 2015) and stabilizing Mad2 protein levels (Nelson *et al*., 2015), CMT-1 is also required to localize PCH-2 to unattached kinetochores during the spindle checkpoint response and generate a robust spindle checkpoint response in AB cells (Nelson *et al*., 2015). Overexpressing Mad2 does not suppress the partial defect in spindle checkpoint activation in *cmt-1* mutants (Nelson *et al*., 2015), suggesting that the defect in spindle checkpoint strength is not because of reduced Mad2 protein levels but the inability to localize PCH-2 to unattached kinetochores.

Here, we test this possibility and show that PCH-2 controls spindle checkpoint strength in *C. elegans*. Despite being essential for the spindle checkpoint in the large somatic, or AB, cell of the 2-cell embryo (Nelson *et al*., 2015), PCH-2 becomes dispensable for the spindle checkpoint and partially dispensable for Mad2 recruitment at unattached kinetochores as AB cells are genetically manipulated to become smaller. The requirement for PCH-2 in promoting spindle checkpoint strength is also observed as cells decrease in size during embryogenesis and in germline precursor, or P_1_, cells, which have a stronger checkpoint than their somatic counterparts. PCH-2 is enriched in P_1_ cells and this enrichment depends on conserved regulators of embryonic polarity, PAR-1 and PAR-6. Further, the stronger checkpoint in P_1_ cells also relies on the *C. elegans* ortholog of p31^comet^, CMT-1, indicating that CMT-1’s ability to enrich PCH-2 in P_1_ cells, in addition to its role in localizing PCH-2 to unattached kinetochores, contributes to a stronger checkpoint. We propose that PCH-2, and its mammalian ortholog TRIP13, ensure a robust spindle checkpoint response and proper chromosome segregation by regulating the availability of O-Mad2 at and near unattached kinetochores. This role may be specifically relevant in scenarios where maintaining genomic stability is particularly challenging, such as in oocytes and early embryos enlarged for developmental competence and germline cells that maintain immortality.

## RESULTS

### PCH-2 becomes dispensable for the spindle checkpoint response in somatic cells experimentally reduced in size

In the large somatic, or AB, cell of the *C. elegans* 2-cell embryo, PCH-2 is essential for spindle checkpoint activation (Nelson *et al*., 2015). To further assess the requirements for PCH-2 function, we manipulated the cell volume of embryos, and thus AB cells, experimentally by performing RNA interference (RNAi) against *ani-2. ani-2* encodes a germline specific anillin whose depletion generates oocytes and, after fertilization, embryos, of varying size (Maddox *et al*., 2005) (Figure 1A). We monitored the length of mitosis in these AB cells, using the time between nuclear envelope breakdown (NEBD) to the onset of cortical contractility (OCC) as markers for the entry into and exit from mitosis, respectively (Essex *et al*., 2009). We then correlated the length of mitosis to cytoplasmic volume. RNAi of *ani-2* did not affect cell cycle progression in control, *pch-2*, or *mad-1* mutants (Figure S1A), indicating that reducing cytoplasmic volume did not affect mitotic timing in AB cells. (In *C. elegans*, the genes that encode Mad1 and Mad2 are *mdf-1* and *mdf-2*, respectively. To avoid confusion, we will use *mad-1* and *mad-2*).

**Figure 1:**
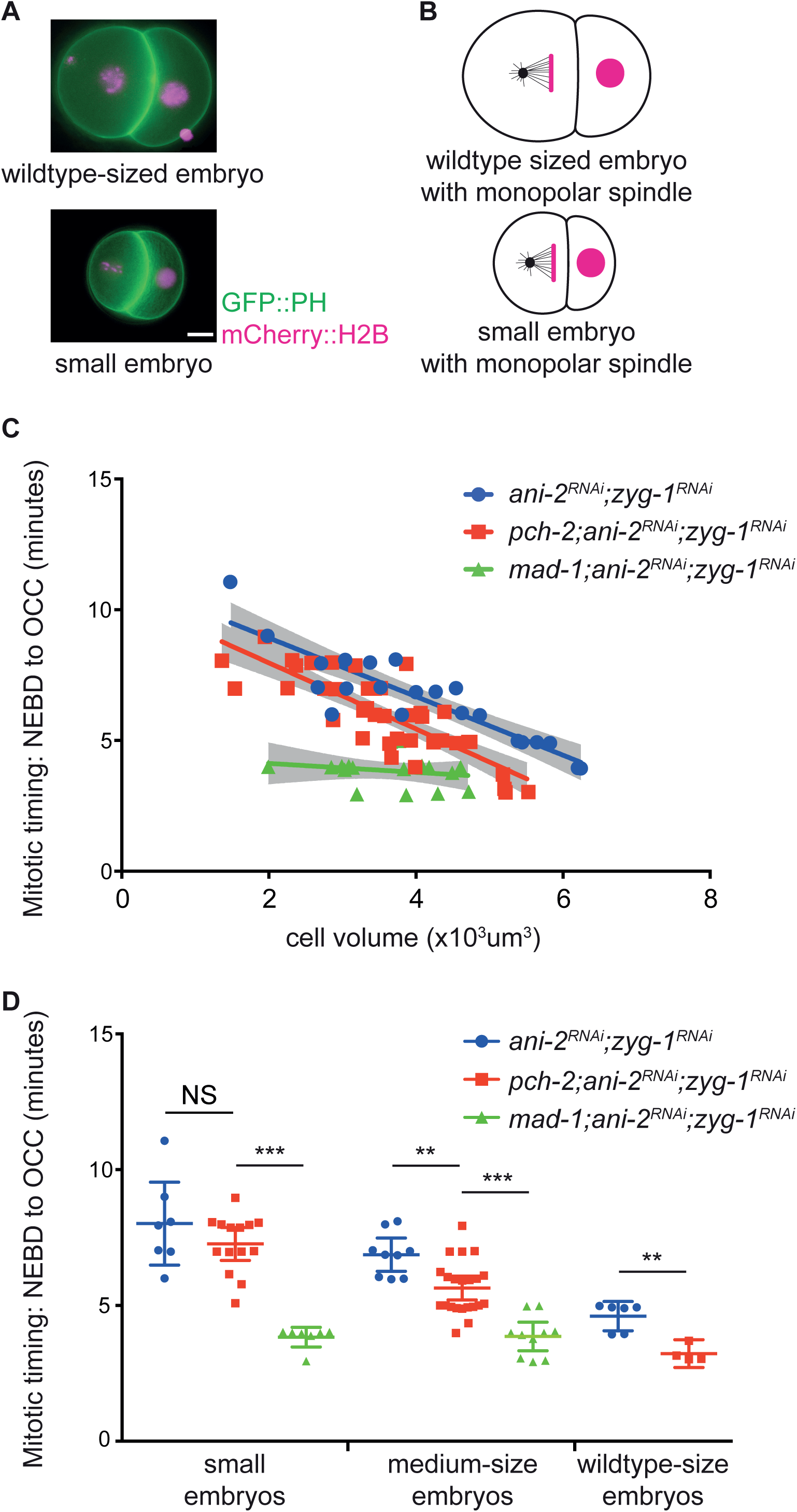
PCH-2 becomes dispensable for the spindle checkpoint response in somatic cells experimentally reduced in size. (A) Images of wildtype-sized and small *ani-2*^*RNAi*^ 2-cell embryos. Scale bar indicates 5 μm. (B) Cartoon of wildtype-sized and small *ani-2*^*RNAi*^ 2-cell embryos treated with *zyg-1*^*RNAi*^. (C) Mitotic timing, as measured from nuclear envelope breakdown (NEBD) to the onset of cortical contractility (OCC), in AB cells of control, *pch-2* and *mad-1* mutant embryos plotted against cell volume. Lines represent least-squares regression models with 95% confidence intervals (gray shaded areas) for each set of data. Equations and p values indicating whether slopes are significantly non-zero for each model are: *ani-2*^*RNAi*^;*zyg-1*^*RNAi*^ (blue): y=-1.117x+11.15 and p < 0.0001; *pch-2;ani-2*^*RNAi*^;*zyg-1*^*RNAi*^ (red): y=-1.264x+10.50 and p < 0.0001; *mad-1;ani-2*^*RNAi*^;*zyg-1*^*RNAi*^ (green): y=-0.1709x+4.468 and p = 0.4197. (D) Data from (C) partitioned into three categories: wild-type sized embryos (more than 5 x 10^3^ μm^3^), medium sized embryos (between 3.3 x 10^3^ μm^3^ and 5 x 10^3^ μm^3^) and small embryos (less than 3.3 x 10^3^ μm^3^). Error bars are 95% confidence intervals. In all graphs, a * indicates a p value < 0.05, ** indicates a p value < 0.01 and *** a p value < 0.0001.

We performed double depletion of *ani-2* and *zyg-1* to induce the spindle checkpoint response in control embryos, *pch-2*, and *mad-1* mutants. ZYG-1 is essential for centrosome duplication and after the first embryonic division, its depletion generates monopolar spindles (O’Connell *et al*., 2001) and unattached kinetochores (Essex *et al*., 2009) (Figure 1B). Consistent with previous reports, as AB cells decreased in cell volume the length of the cell cycle delay, an indicator of spindle checkpoint strength, increased in control embryos (Galli and Morgan, 2016; Gerhold *et al*., 2018) (Figure 1C, Videos 1 and 2). Surprisingly, as *pch-2* mutants decreased in size, the spindle checkpoint response more closely resembled that of control AB cells than *mad-1* mutants (Figure 1C, Videos 3 and 4). *mad-1* mutant embryos appear more sensitive to *ani-2* RNAi treatment and we had difficulty recovering any wild-type sized *mad-1* embryos. There was no significant difference between the slopes of the regression analysis of control and *pch-2* mutant data (p value = 0.4664), while the slopes between the regression analysis of *pch-2* and *mad-1* mutant data were significantly different (p value = 0.0007). To make these comparisons more clear, we binned our data. By our measurements, control AB cells ranged from 5 to 6 × 10^3^ μm^3^. Therefore, we classified AB cells which were more than 5 × 10^3^ μm^3^ as wildtype sized. The remaining cells, which ranged from 1.5 × 10^3^ μm^3^ to 5 × 10^3^ μm^3^ were partitioned equally into two classes: medium sized embryos were between 3.3 × 10^3^ μm^3^ and 5 × 10^3^ μm^3^ and small embryos were between 1.5 × 10^3^ μm^3^ and 3.3 × 10^3^ μm^3^. When partitioned into these classes, *pch-2* mutants produced no checkpoint response in wildtype-sized AB cells, as previously reported (Nelson *et al*., 2015), defects in checkpoint strength in medium-sized AB cells and a robust checkpoint response in small AB cells (Figure 1D).

We verified that the mitotic delay observed in *pch-2* AB cells was a legitimate spindle checkpoint response by monitoring mitotic timing after performing double depletion of *ani-2* and *zyg-1* in *san-1* and *pch-2;san-1* mutant embryos. SAN-1 is the *C. elegans* ortholog of the essential spindle checkpoint factor, Mad3 (Nystul *et al*., 2003) (Figure S1B). There was no significant difference between the slopes of the regression analysis of *san-1* and *pch-2;san-1* data (p value = 0.8813) and the slopes of each model were not statistically different than zero (Figure S1B). However, we observed a slight increase in the length of the cell cycle as cells got smaller in *san-1* mutants, potentially reflecting that the spindle checkpoint in *C. elegans* is composed of two independent branches (Essex *et al*., 2009). These data allow us to draw two important conclusions: 1) Since we observe robust spindle checkpoint activation in *pch-2* mutant AB cells as they decrease in size and mitotic timing is similar to what we observe in control cells, PCH-2 does not appear to affect spindle checkpoint silencing in *C. elegans*; and 2) the requirement for PCH-2 during spindle checkpoint activation is proportional to cell volume in AB cells with monopolar spindles.

### MAD-2 recruitment is partially restored to unattached kinetochores in *pch-2* somatic cells experimentally reduced in size

We showed that PCH-2 is required for robust recruitment of Mad2 at unattached kinetochores during spindle checkpoint activation in AB cells of 2-cell embryos (Nelson *et al*., 2015). Therefore, we tested whether the checkpoint induced delay we observed in small *ani-2*^*RNAi*^;*zyg-1*^*RNAi*^;*pch-2* AB cells was accompanied by increased recruitment of GFP::MAD-2 at unattached kinetochores. We quantified GFP::MAD-2 recruitment at unattached kinetochores in pseudo-metaphase in control animals and *pch-2* mutants treated with *ani-2* and *zyg-1* RNAi (Figure 2A) and plotted GFP::MAD-2 fluorescence against cell volume (Figure 2B). Surprisingly, the regression analysis for control AB cells had a positive slope, suggesting that less GFP::MAD-2 is required at unattached kinetochores for spindle checkpoint function as these cells became smaller (Figure 2B). This was despite similar levels of soluble GFP::MAD2 around mitotic chromosomes after NEBD in both genetic backgrounds (Figures S2A and S2B). We observed that the regression analysis of GFP::MAD-2 fluorescence at unattached kinetochores in *pch-2;ani-2*^*RNAi*^;*zyg-1*^*RNAi*^ AB cells exhibited a negative slope, showing improved GFP::MAD-2 recruitment to unattached kinetochores as cells got smaller. However, the amount of GFP::MAD-2 was typically lower in fluorescence intensity than *ani-2*^*RNAi*^;*zyg-1*^*RNAi*^ control cells (Figure 2B). Therefore, our experiments demonstrate that MAD-2 recruitment is partially restored to unattached kinetochores in *pch-2* mutant somatic cells experimentally reduced in size.

**Figure 2:**
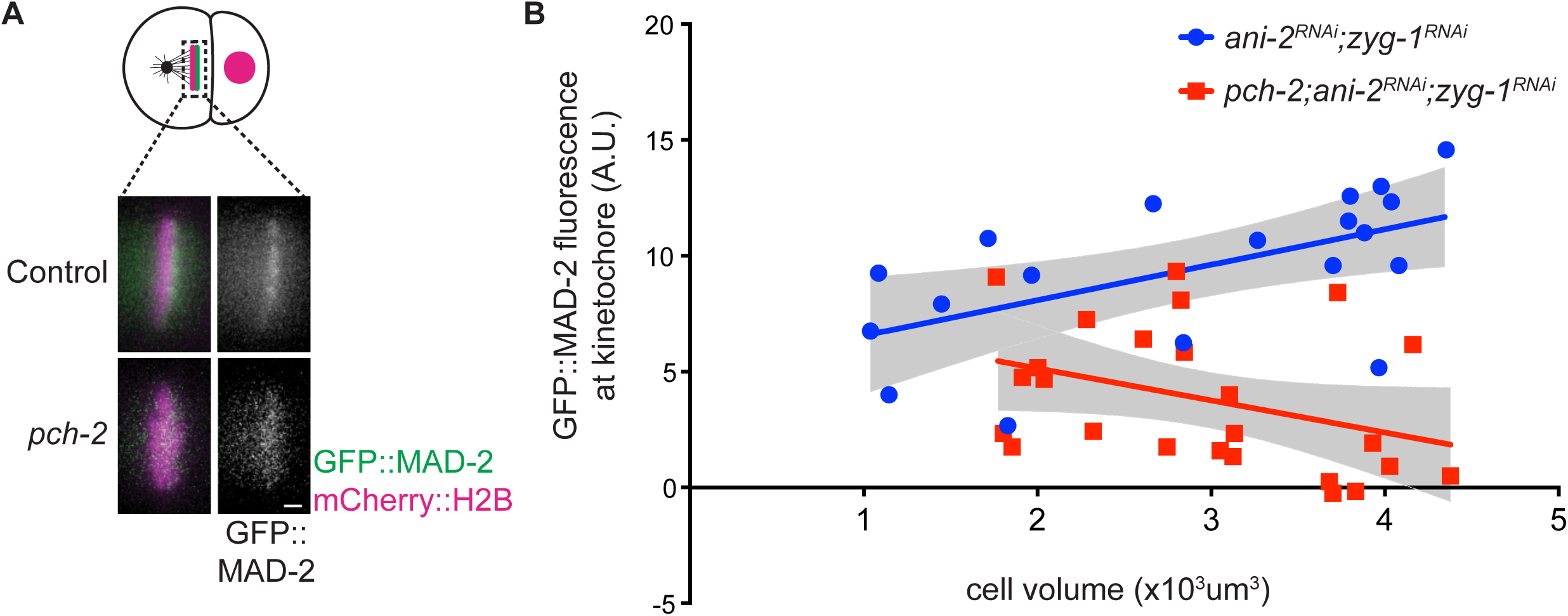
MAD-2 recruitment is partially restored to unattached kinetochores in *pch-2* mutant somatic cells experimentally reduced in size. (A) Cartoon and images of GFP::MAD-2 recruitment to unattached kinetochores in AB cells of control and *pch-2* AB cells treated with *ani-2* and *zyg-1* RNAi. Scale bar indicates 1 μm. (B) Quantification of kinetochore bound GFP::MAD-2 in control and *pch-2* AB cells plotted against cell volume. Lines represent least-squares regression models with 95% confidence intervals (gray shaded areas) for each set of data. Equations and p values indicating whether slopes are significantly non-zero for each model are: *ani-2*^*RNAi*^;*zyg-1*^*RNAi*^ (blue): y=1.531x+5.024 and p = 0.0115; *pch-2;ani-2*^*RNAi*^;*zyg-1*^*RNAi*^ (red): y=-1.384x+7.911 and p = 0.0384.

### MAD-2 dosage controls checkpoint strength

*C. elegans* meiotic nuclei in the germline exist in a syncytium and cellularize after completing meiotic prophase. Knockdown of *ani-2* affects this cellularization event, resulting in a loss of cytoplasmic volume after nuclei are fully formed (Maddox *et al*., 2005). MAD-2 is localized to the nucleus and nuclear envelope in these oocytes (Bohr *et al*., 2015; Lawrence *et al*., 2015) (Figure 3A). Since embryonic nuclear size is not affected by *ani-2* RNAi (Figure S3), we reasoned that as cells are genetically manipulated to decrease in cell volume, the absolute amount of Mad2 protein is likely to remain constant but its concentration increases. Given that TRIP13 function is dispensable for checkpoint activation when O-Mad2 is readily available in mammalian cells (Ma and Poon, 2018), we reasoned that something similar might be happening in *C. elegans* embryos. Specifically, we hypothesized that an increase in concentration of Mad2, and O-Mad2 in particular, may explain the reduced requirement for PCH-2 in *ani-2*^*RNAi*^;*zyg-1*^*RNAi*^ small AB cells (Figure 3B).

**Figure 3:**
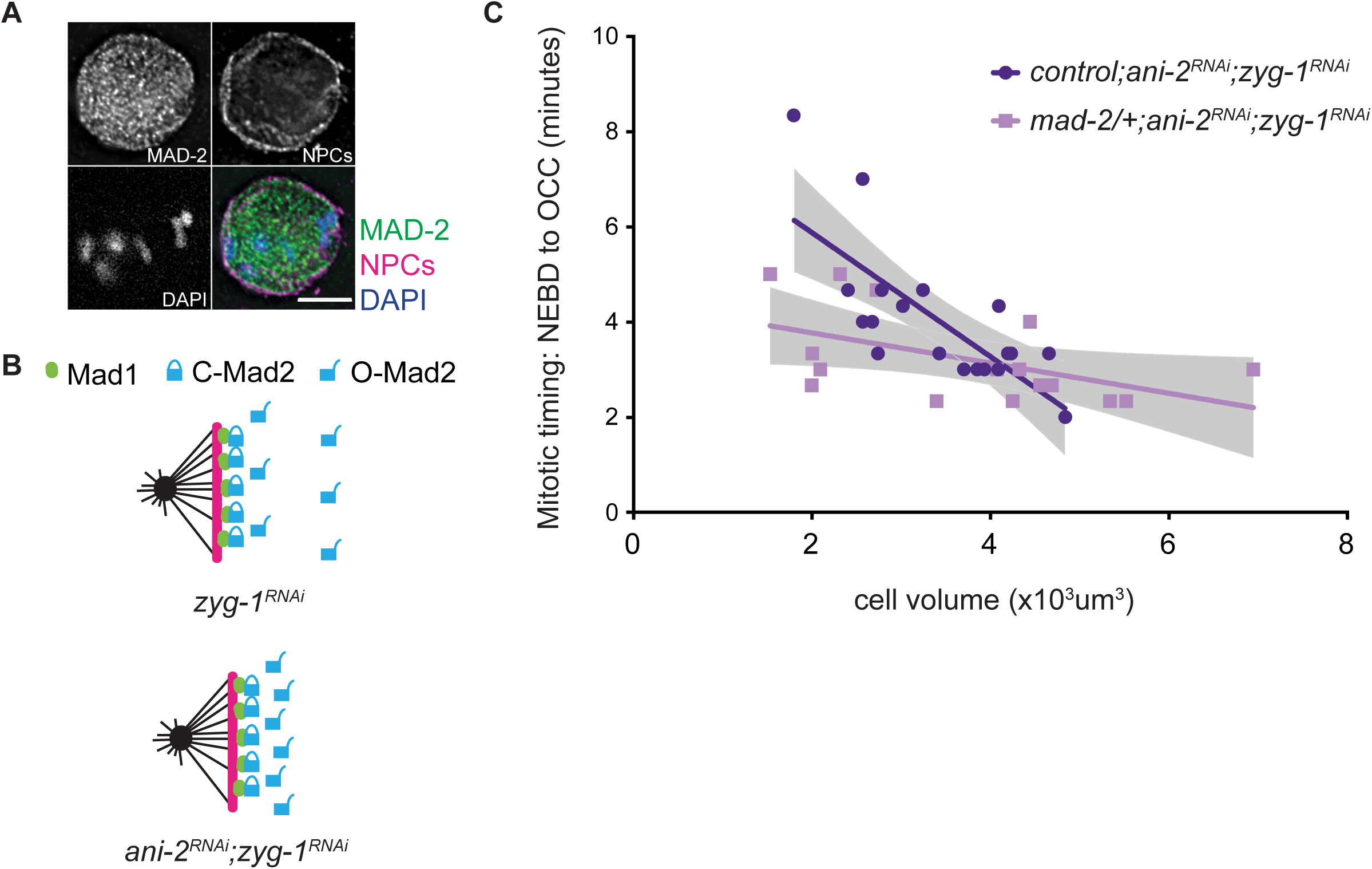
MAD-2 dosage controls checkpoint strength. (A) Immunostaining of MAD-2 and nuclear pore complex components (NPCs) shows MAD-2 localized in the nucleus and at the nuclear envelope during interphase. Scale bar indicates 5 μm. (B) Model depicting how a decrease in cell volume might result in an increase in the local concentration of O-Mad2 in *ani-2*^*RNAi*^;*zyg-1*^*RNAi*^ embryos, in contrast to *zyg-1*^*RNAi*^ embryos. (C) Mitotic timing, as measured from nuclear envelope breakdown (NEBD) to the onset of cortical contractility (OCC), in AB cells of control and *mad-2/+* mutant embryos plotted against cell volume. Lines represent least-squares regression models with 95% confidence intervals (gray shaded areas) for each set of data. Equations and p values indicating whether slopes are significantly non-zero for each model are: control (dark purple): y=-1.302x+8.477 and p = 0.0002; *mad-2/+* (light purple): y=-0.3171x+4.402 and p = 0.0395.

To test this possibility, we initially attempted to directly visualize O-Mad2 in *C. elegans* embryos. Unfortunately, we were unable to perform this experiment with a commercial antibody (data not shown). Further, we could not directly probe total Mad2 concentration as cells decrease in volume upon treatment with *ani-2* RNAi because GFP::MAD-2 does not localize to the nucleus and instead localizes in the cytoplasm until NEBD (Essex *et al*., 2009; Nelson *et al*., 2015), making it an inaccurate reporter for this assay. Instead, we tested whether reducing Mad2 dosage affected checkpoint strength. We hypothesized that if Mad2 concentration influences checkpoint strength, reducing it by half should attenuate checkpoint strength in comparison to control animals. We performed double depletion of *ani-2* and *zyg-1* by RNAi in *mad-2* heterozygotes. Indeed, *mad-2* heterozygotes exhibited stronger spindle checkpoint strength as cells became smaller. However, the increase in spindle checkpoint strength was less robust than control cells (Figure 3C). The slopes of the linear regressions for both control and *mad-2* heterozygotes were significantly non-zero, unlike similar experiments with *mad-1* and *san-1* homozygotes (Figures 1B and S1D). Therefore, spindle checkpoint strength depends on MAD-2 dosage.

We wondered whether the decrease in Mad2 protein levels might restore the reliance on PCH-2 in small embryos. However, *pch-2;mad-2/+* double mutants exhibited a substantial decrease in the production and viability of embryos, preventing us from performing these experiments: *pch-2;mad-2/+* double mutants produced broods that were 14% of control animals and only 1% of these embryos were viable. Further, *pch-2;mad-2* double mutants could not be recovered from *pch-2;mad-2/+* mothers, a genetic interaction that we did not observe when we generated *pch-2;mad-1* double mutants (Bohr *et al*., 2015) or *pch-2;san-1* double mutants (Figure S1B). Worms with mutations in some spindle checkpoint mutants often display defects in fertility, viability and development (Kitagawa and Rose, 1999; Stein *et al*., 2007; Lara-Gonzalez *et al*., 2019). Thus, in addition to MAD-2 dosage controlling checkpoint strength, it collaborates with PCH-2 to promote *C. elegans* fertility and viability.

### PCH-2 affects spindle checkpoint strength during embryogenesis

During embryogenesis, cell volume decreases and spindle checkpoint strength increases (Galli and Morgan, 2016; Gerhold *et al*., 2018). Given that the requirement for PCH-2 is proportional to cell volume in 2-cell embryos treated with *ani-2* RNAi, we assessed the role for PCH-2 in spindle checkpoint activation as cells decreased in size during normal embryogenesis.

We permeabilized embryos by performing *perm-1* RNAi (Carvalho *et al*., 2011) and activated the spindle checkpoint by treating these *perm-1*^*RNAi*^ embryos with nocodazole, a drug that depolymerizes microtubules. Since we could not reliably visualize OCC in these dividing embryos, we measured mitotic timing from nuclear envelope break down (NEBD) to decondensation of chromosomes (DECON) in cells of the AB lineage. These cells in control embryos exhibited a longer mitotic delay in 16-cell than in 4-cell embryos (Figure 4A), verifying that the spindle checkpoint increases in strength as cells decrease in volume during embryogenesis (Galli and Morgan, 2016; Gerhold *et al*., 2018). As a control, we performed the same experiment in *san-1* mutants and did not detect a mitotic delay when these embryos were treated with nocodazole (Figure 4A). Cells in 4-cell *pch-2* mutant embryos treated with nocodazole resembled *san-1* mutants (Figure 4A). However, cells in the AB lineage in 16-cell *pch-2* mutant embryos treated with nocodazole exhibited a significant cell cycle delay when compared to similar cells in *pch-2* mutants treated with DMSO and *san-1* mutants treated with nocodazole. These delays were not as prolonged as that observed in control embryos treated with nocodazole (Figure 4A). Thus, as cells of the AB lineage naturally decrease in cell size to 16 cell embryos, *pch-2* mutants treated with nocodazole delay the cell cycle but exhibit defects in spindle checkpoint strength.

Given our hypothesis that Mad2 dosage might contribute to spindle checkpoint strength, particularly in *pch-2* mutants, we tested if a subtle increase in MAD-2 protein levels would suppress the defect in spindle checkpoint function or strength in *pch-2* mutant embryos. The presence of a GFP::MAD-2 transgene, in addition to endogenous MAD-2, results in about 2.5 times more MAD-2 in worms. This slight overexpression generates a normal spindle checkpoint response in control AB cells and can bypass the requirement for checkpoint components MAD-3 or BUB-3 (Essex et al., 2009), but not PCH-2 (Nelson et al., 2015) in AB cells of 2-cell embryos with monopolar spindles. Overexpression of MAD-2 did not affect the checkpoint response in 16-cell *pch-2* embryos (Figure 4B). However, in contrast to our results in 4-cell *pch-2* mutant embryos treated with nocodazole (Figure 4A), we found that overexpression of MAD-2 in cells of the AB lineage of 4-cell *pch-2* embryos produced cell cycle delays when compared to the same cells in embryos treated with DMSO. Again, these delays were not as dramatic as control cells overexpressing GFP::MAD-2 (Figure 4B) but were significant, allowing us to conclude that slight overexpression of Mad2 partially restores checkpoint function to *pch-2* mutants as cells of the AB lineage decrease in size during embryogenesis, at least in 4-cell embryos.

**Figure 4:**
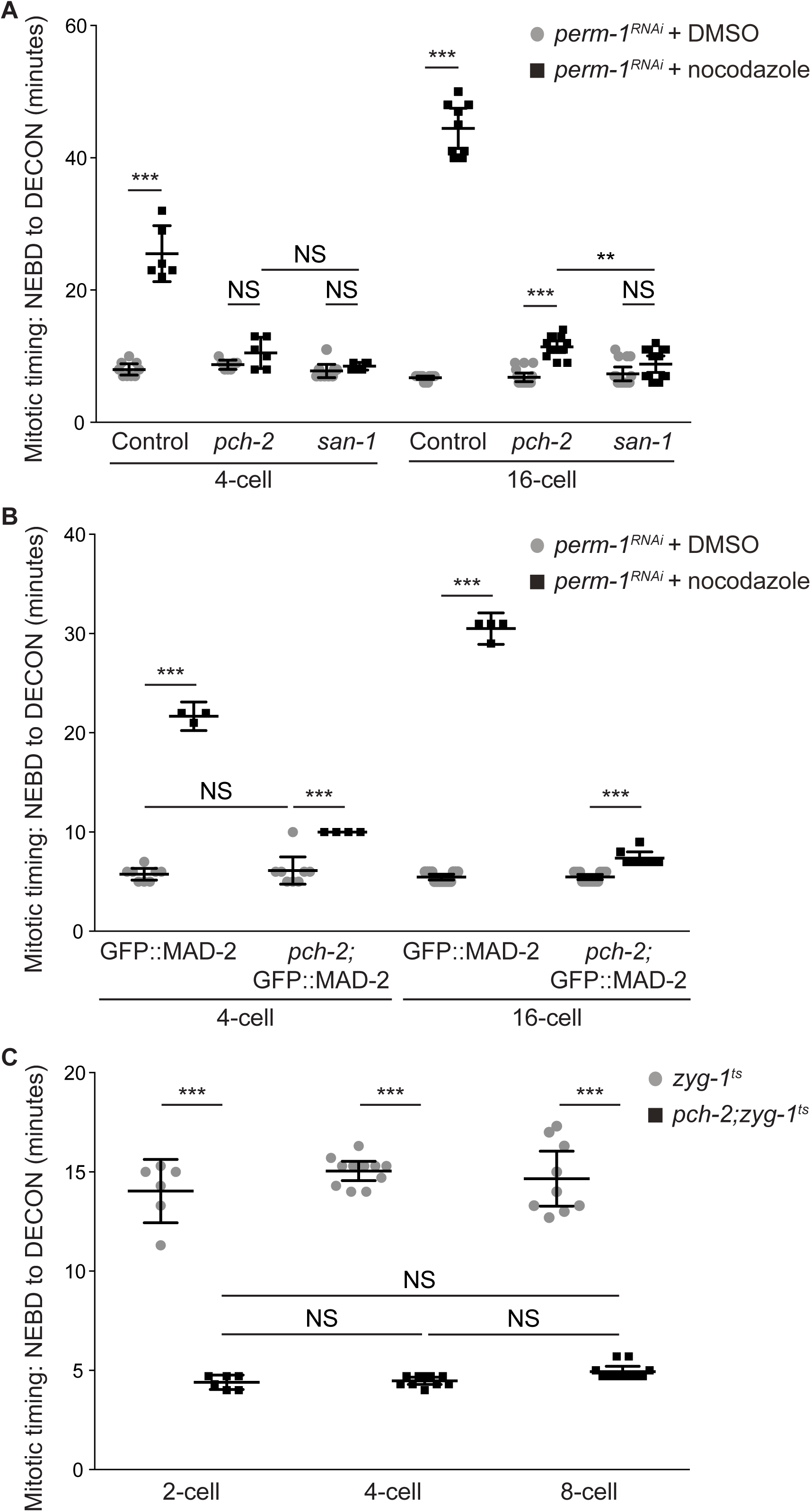
PCH-2 regulates spindle checkpoint strength during embryogenesis. (A) Mitotic timing, as measured from nuclear envelope breakdown (NEBD) to decondensation of chromatin (DECON), in control, *pch-2* and *san-1* mutant embryos treated with *perm-1* RNAi and DMSO or nocodazole at different developmental stages (4- and 16-cell embryos). (B) Mitotic timing in control and *pch-2* mutant embryos overexpressing GFP::MAD-2 and treated with *perm-1* RNAi and either DMSO or nocodazole at different developmental stages (4- and 16-cell embryos). (C) Mitotic timing in *zyg-1*^*ts*^ and *pch-2;zyg-1*^*ts*^ mutant embryos at different developmental stages (2-, 4- and 8-cell embryos). All error bars are 95% confidence intervals.

Given that we activated the spindle checkpoint in *ani-2*^*RNAi*^ embryos by generating monopolar spindles (Figures 1C and 3C), we also tested whether embryos with monopolar spindles might behave differently than embryos treated with nocodazole, particularly in very early embryogenesis. We used a fast acting temperature sensitive allele of *zyg-1* (*zyg-1*^*ts*^) (O’Rourke *et al*., 2011) to activate the spindle checkpoint in developing embryos with 2, 4 and 8 cells. We shifted embryos at different stages of development, verified the appearance of monopolar spindles and measured mitotic timing from NEBD to DECON. In control *zyg-1*^*ts*^ mutant embryos, we observed a delay in mitotic timing in cells from the AB lineage and this delay only became marginally longer as embryos had more cells (Figure 4C), similar to previous reports (Gerhold *et al*., 2018). In stark contrast to our *ani-2*^*RNAi*^ experiments, the mitotic timing observed in *pch-2;zyg-1*^*ts*^ mutant embryos was the same in AB cells of 2-cell, 4-cell and 8-cell embryos and significantly reduced in comparison to *zyg-1*^*ts*^ embryos. Thus, similar to our results with 4-cell *pch-2* mutant embryos treated with nocodazole, *pch-2* mutants exhibit no cell cycle delay in the presence of monopolar spindles in AB cells in 2-cell, 4-cell and 8-cell embryos. However, additional considerations may make direct comparisons between our *ani-2*^*RNAi*^;*zyg-1*^*RNAi*^ experiments and *zyg-1*^*ts*^ embryos difficult (see Discussion).

### PCH-2 is responsible for the stronger spindle checkpoint in the germline lineage

Cell fate is another important determinant of spindle checkpoint strength. In *C. elegans* embryos, the spindle checkpoint is stronger in germline precursor cells than similarly sized somatic counterparts (Galli and Morgan, 2016; Gerhold *et al*., 2018). However, as we observed with AB cells (Nelson *et al*., 2015), PCH-2 is essential for the spindle checkpoint in wildtype-sized P_1_ cells (Figure S4). Therefore, having established that PCH-2 becomes dispensable for the spindle checkpoint as 2-cell embryos are genetically manipulated to become smaller (Figures 1C and D), we tested whether PCH-2 contributed to the stronger spindle checkpoint in P_1_ cells of 2-cell embryos treated with *ani-2* RNAi (Figures 5 and S4B). Consistent with other reports (Galli and Morgan, 2016; Gerhold *et al*., 2018), when we performed double depletion of *ani-2* and *zyg-1* in control embryos and monitored mitotic timing, we observed P_1_ cells with similar volumes as AB cells exhibiting a longer cell cycle delay (Figures 5A and S4B, Videos 5 and 6). Further, the regression analysis that best fit control P_1_ data is significantly different and steeper than that of control AB cells (p value < 0.0001), indicating that variables in addition to cell volume contribute to the spindle checkpoint strength in germline precursor cells. When we knocked down both *ani-2* and *zyg-1* in *pch-2* mutant embryos, we no longer observed a significant difference (p value = 0.9096) between the slopes of the regression analysis of P_1_ and AB cells (Figures 5B and S4B, Videos 7 and 8), indicating that PCH-2 is responsible for the stronger checkpoint in P_1_ cells.

**Figure 5:**
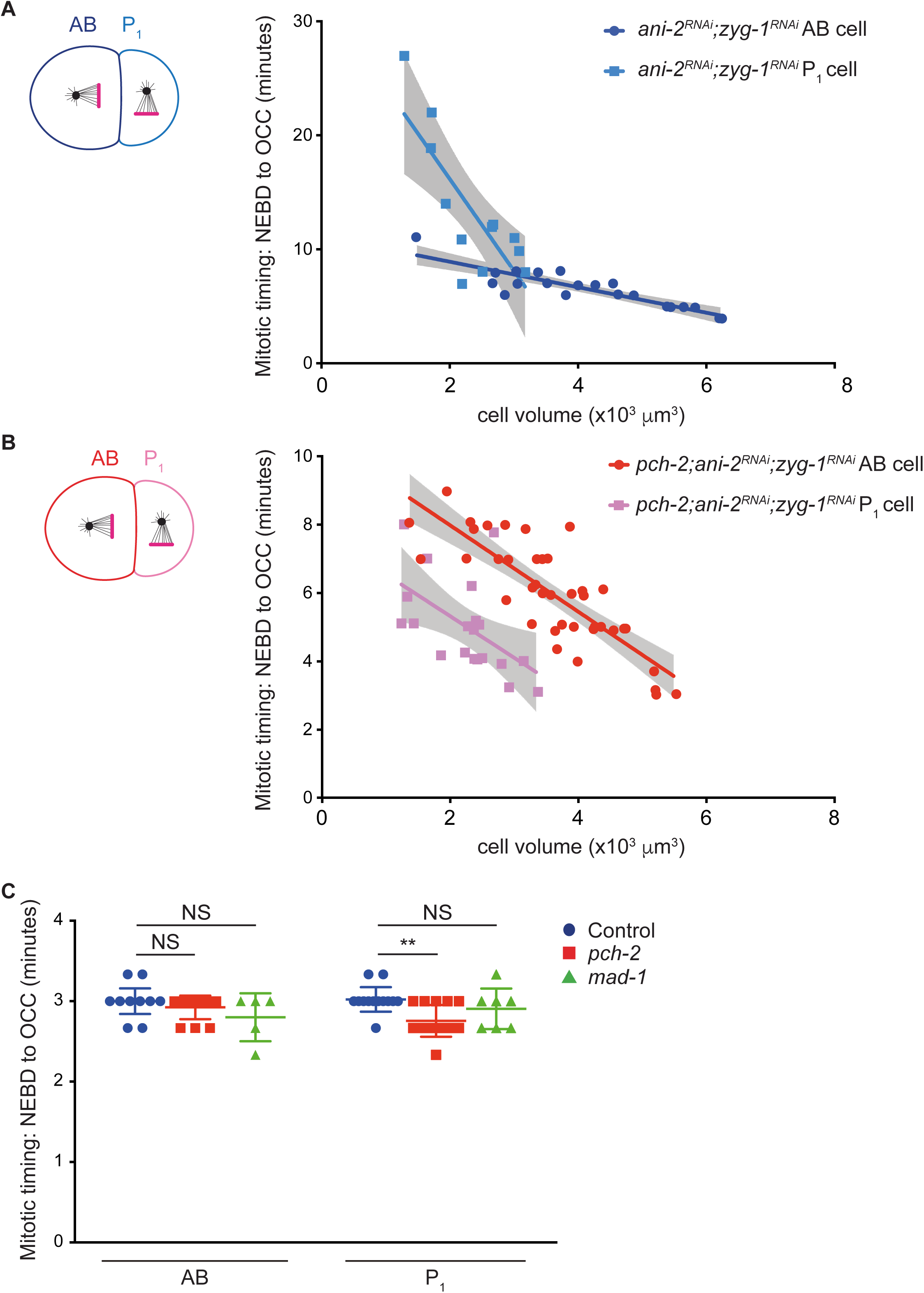
PCH-2 is responsible for the stronger spindle checkpoint in the germline lineage. Mitotic timing, as measured from nuclear envelope breakdown (NEBD) to the onset of cortical contractility (OCC), in AB and P_1_ cells plotted against cell volume in control *ani-2*^*RNAi*^;*zyg-1*^*RNAi*^ embryos (A) or *pch-2;ani-2*^*RNAi*^;*zyg-1*^*RNAi*^ (B) embryos. Lines represent least-squares regression models with 95% confidence intervals (gray shaded areas) for each set of data. Equations and p values indicating whether slopes are significantly non-zero for each model are: *ani-2*^*RNAi*^;*zyg-1*^*RNAi*^ AB (dark blue): y=-1.117x+11.15 and p < 0.0001; *ani-2*^*RNAi*^;*zyg-1*^*RNAi*^ P_1_ (light blue): y=-8.047x+32.27 and p = 0.0021; *pch-2;ani-2*^*RNAi*^;*zyg-1*^*RNAi*^ AB (red): y=-1.264x+10.50 and p < 0.0001; *pch-2;ani-2*^*RNAi*^;*zyg-1*^*RNAi*^ P_1_ (pink): y=-1.218x+7.75 and p = 0.0125. Data for AB cells in both control and *pch-2* mutants is the same as in Figure 1C. (C) Mitotic timing of AB and P_1_ cells in control, *pch-2* and *mad-1* mutants during unperturbed divisions. Error bars are 95% confidence intervals.

We observed that cell cycle timing was faster in *pch-2* mutant P_1_ cells than similarly sized *pch-2* mutant AB cells after treatment with *ani-2* and *zyg-1* RNAi (Figures 5B and S4B). We wondered if embryonic germline precursor cells might rely on some spindle checkpoint proteins for normal mitotic timing, analogous to mitotically dividing stem cells in the *C. elegans* germline (Gerhold *et al*., 2015) and similar to mammalian cultured cells (Meraldi *et al*., 2004; Rodriguez-Bravo *et al*., 2014; Ma and Poon, 2016). To address this, we measured normal mitotic timing in AB and P_1_ cells of both control and *pch-2* mutant embryos. We found that while normal mitotic timing is unaffected by mutation of *pch-2* in AB cells, *pch-2* mutant P_1_ cells go through mitosis significantly faster than control P_1_ cells (Figure 5C), thus providing an explanation for the faster cell cycle timing in *pch-2* mutant P_1_ cells with the same cell volume as *pch-2* mutant AB cells after treatment with *ani-2* and *zyg-1* RNAi. We saw a decrease in the cell cycle timing of P_1_ cells in *mad-1* mutants but this was not significantly different than control P_1_ cells (Figure 5C).

### PCH-2’s enrichment in P_1_ cells depends on PAR-1 and PAR-6

Cell fate is driven by the asymmetric distribution of various determinants between somatic and germline lineages during early divisions of the *C. elegans* embryo (Rose and Gonczy, 2014). Since we found that PCH-2 promoted the spindle checkpoint strength in both AB and P_1_ cells, but even more dramatically in P_1_ cells, we asked if PCH-2 was regulated differently between these cells. First, we tested whether PCH-2::GFP could also support the stronger checkpoint in P_1_ cells. We treated embryos expressing PCH-2::GFP with *zyg-1* RNAi and evaluated mitotic timing in both AB and P_1_ cells, in control cells and in the presence of monopolar spindles, using chromosome decondensation as a marker for mitotic exit. P_1_ cells expressing PCH-2::GFP had full checkpoint function, exhibiting a mitotic delay longer than AB cells also expressing PCH-2::GFP (Figure 6A).

**Figure 6:**
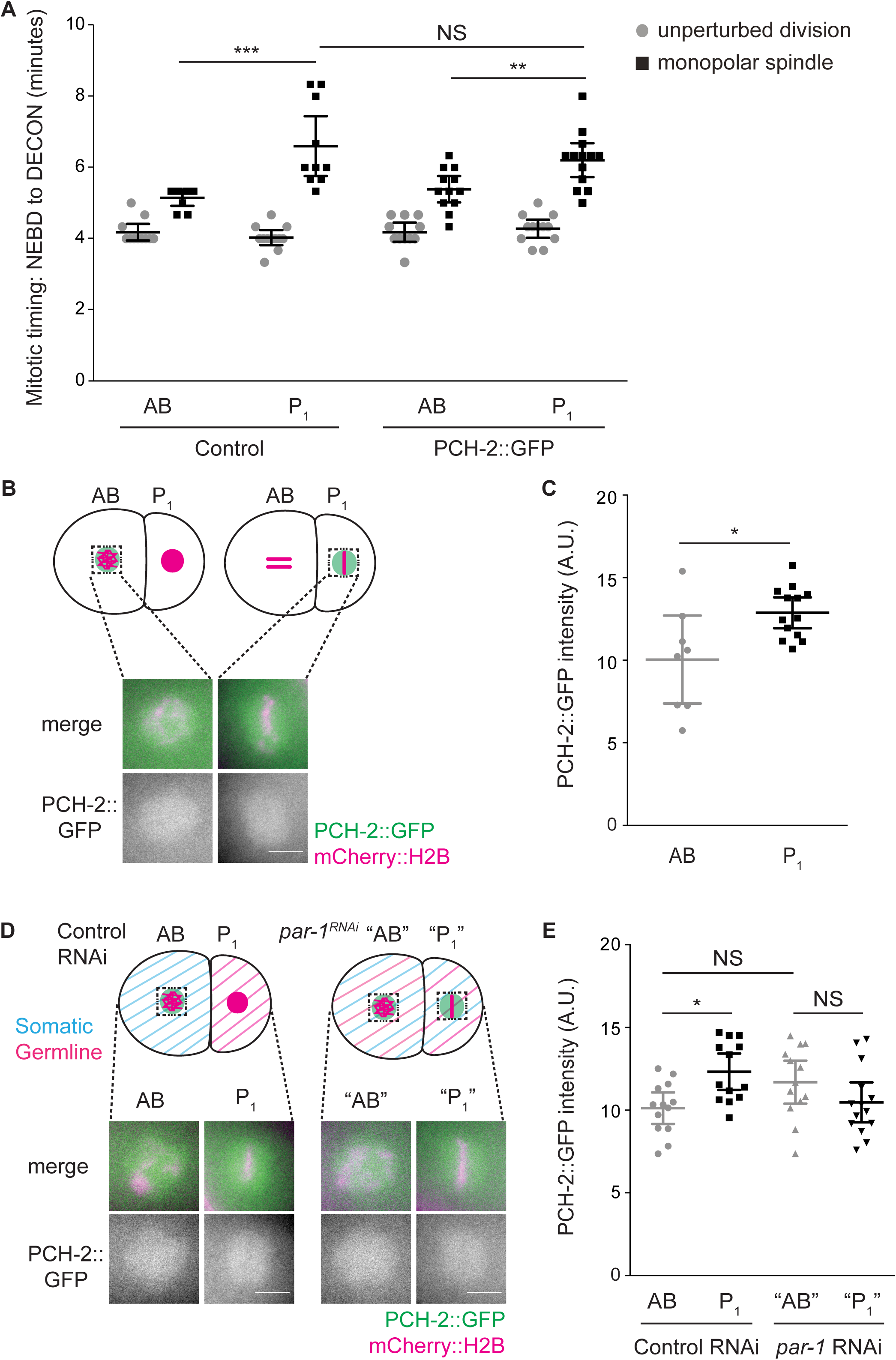
PCH-2’s enrichment around mitotic chromosomes in P_1_ cells depends on PAR-1. (A) Mitotic timing of control embryos and embryos expressing PCH-2::GFP during unperturbed divisions or in the presence of monopolar spindles. (B) Cartoon and images of PCH-2::GFP localization around mitotic chromosomes in AB and P_1_ cells of 2-cell embryos. Scale bar indicates 5 μm. (C) Quantification of PCH-2::GFP fluorescence in AB and P_1_ cells. (D) Cartoon and images of PCH-2::GFP localization around mitotic chromosomes in AB and P_1_ cells of control RNAi and *par-1*^*RNAi*^ 2-cell embryos. (E) Quantification of PCH-2::GFP fluorescence in AB and P_1_ cells of *par-1*^*RNAi*^ embryos. All error bars are 95% confidence intervals. NS indicates not significant.

Previous transcriptome analysis of PCH-2 did not reveal asymmetric enrichment of PCH-2 mRNA between AB and P_1_ cells (Tintori *et al*., 2016). We tested whether PCH-2::GFP exhibited differences in protein levels between AB and P_1_ cells. First, we assessed whether PCH-2::GFP was more enriched in pseudo-metaphase at unattached kinetochores in P_1_ than AB cells. We quantified PCH-2::GFP fluorescence at unattached kinetochores in both AB and P_1_ cells of embryos treated with *zyg-1* RNAi but did not detect any difference between the two cell types (Figures S5A and B). Similarly, we did not detect any difference in GFP::MAD-2 recruitment at unattached kinetochores between AB and P_1_ cells in *zyg-1*^*RNAi*^ embryos (Figures S5C and D).

Checkpoint factors, including MAD-2 and PCH-2, form a diffuse “cloud” around mitotic chromosomes after NEBD, even during normal cell cycles (Essex *et al*., 2009; Nelson *et al*., 2015). We wondered if PCH-2::GFP fluorescence in this cloud might be different between AB and P_1_ cells. First, we verified that PCH-2::GFP fluorescence around mitotic chromosomes was similar between AB cells during unperturbed (control) or monopolar mitosis (*zyg-1*^*RNAi*^) (Figure S6). PCH-2::GFP fluorescence around chromosomes was significantly higher in *zyg-1*^*RNAi*^ AB cells than control AB cells (Figure S6B). However, we noticed that the area occupied by PCH-2::GFP in control AB cells was significantly larger than that of *zyg-1*^*RNAi*^ AB cells (yellow dashed circle in Figure S6A and quantified in Figure S6C). When we factored this larger area of PCH-2::GFP fluorescence into our analysis, we observed a similar amount of PCH-2::GFP around mitotic chromosomes in both control and *zyg-1*^*RNAi*^ AB cells (Figure S6D).

Having established that AB cells had similar amounts of PCH-2::GFP whether the checkpoint was active or not, we quantified PCH-2::GFP fluorescence in the area around mitotic chromosomes in AB and P_1_ cells during unperturbed cell cycles. Similar to AB cells (Nelson *et al*., 2015) and Figure S6A), we observed PCH-2::GFP enriched in the area around the chromosomes in prometaphase in P_1_ cells (Figure 6B). When we quantified the fluorescence of PCH-2::GFP in this area surrounding chromosomes after NEBD in both AB and P_1_ cells, we detected a statistically significant enrichment of PCH-2::GFP in the area surrounding chromosomes in P_1_ cells (Figure 6C) but not in the cytoplasm of P_1_ cells (Figures S7A and B). Although this enrichment is limited to a “cloud” around mitotic chromosomes (see Figure S7B), we verified that this enrichment was not the indirect consequence of the smaller volume of P_1_ cells by quantifying PCH-2::GFP fluorescence in *gpr-1/2*^*RNAi*^ embryos. This double knockdown equalizes the size of AB and P_1_ cells without affecting their cell fate (Colombo *et al*., 2003; Gotta *et al*., 2003; Srinivasan *et al*., 2003). RNAi of *gpr-1/2* showed variability in the effect on AB and P_1_ cell size (Figure S7C). However, when we limited our analysis of PCH-2::GFP fluorescence to embryos in which AB and P_1_ cells were of similar area (red symbols in Figure S7C), we observed a similar enrichment of PCH-2::GFP in P_1_ cells as control embryos (Figures S7D and E).

To better understand the relationship between PCH-2 enrichment in P_1_ cells and cell fate, we abrogated the asymmetry of the 2-cell embryo by performing RNAi against the essential polarity factors, PAR-1 (Guo and Kemphues, 1995) and PAR-6 (Hung and Kemphues, 1999). These factors antagonize each other, with PAR-6 at the anterior cortex and PAR-1 at the posterior cortex of early embryos, to establish asymmetries during the first two embryonic divisions (Goldstein and Macara, 2007). In both *par-1*^*RNAi*^ and *par-6*^*RNAi*^ mutant embryos, AB and P_1_ cells exhibit similar checkpoint strength (Gerhold *et al*., 2018), indicating that the stronger spindle checkpoint response in P_1_ cells depends on this asymmetric division. Despite the loss of cell fate in *par-1*^*RNAi*^ and *par-6*^*RNAi*^ embryos, we will refer to the anterior blastomere as “AB” and the posterior as “P_1_”. We verified the efficiency of *par-1* and *par-6* RNAi by measuring cell area and found that AB and P_1_ cells approached similar sizes in both conditions (Figures S8A and B), although AB cells were still significantly larger than P_1_ cells in *par-1*^*RNAi*^ mutant embryos (Figure S8A). We quantified PCH-2::GFP fluorescence in the area around chromosomes in AB and P_1_ cells after *par-1* RNAi and observed that the fluorescence of PCH-2::GFP, despite being slightly lower in P_1_ cells, was not significantly different between AB and P_1_ cells, unlike what we observed in embryos exposed to control RNAi (Figures 6D and E). AB and P_1_ cells treated with *par-6* RNAi showed equal PCH-2::GFP fluorescence (Figures S8C and D). Therefore, PCH-2::GFP’s enrichment around mitotic chromosomes in P_1_ cells depends on the conserved factors that induce embryonic asymmetry and germline cell fate, PAR-1 and PAR-6.

### The stronger checkpoint in P_1_ cells depends on CMT-1

In vitro, the *C. elegans* ortholog of p31^comet^, CMT-1, is required for PCH-2 to bind and remodel Mad2 (Ye *et al*., 2015). In addition to this role, CMT-1 is also required to localize PCH-2 to unattached kinetochores and generate a robust spindle checkpoint response in AB cells (Nelson *et al*., 2015). Therefore, we reasoned that CMT-1 might also be required for the stronger checkpoint in P_1_ cells.

To test this possibility, we first performed double knockdown of *ani-2* and *zyg-1* in *cmt-1* mutants and monitored the length of the spindle checkpoint response as AB and P_1_ cells became smaller (Figure 7A, Videos 9-12). When compared to the regression model for control AB cells (opaque blue line in Figure 7A), we saw that *cmt-1* AB cells consistently exhibit a weaker checkpoint at all cell volumes. Similar to *pch-2;ani-2*^*RNAi*^;*zyg-1*^*RNAi*^ mutants (Figure 5B), the stronger spindle checkpoint response in P_1_ cells was lost in *cmt-1;ani-2*^*RNAi*^;*zyg-1*^*RNAi*^ mutants and we did not observe any statistical difference between the between the slopes of the regression analysis of P_1_ and AB cells (p value = 0.9403). We also observed that cell cycle timing was faster in *cmt-1* P_1_ cells that were similar in volume to *cmt-1* AB cells (Figure 7A). Thus, CMT-1 is also essential to promote spindle checkpoint strength in germline precursor cells.

**Figure 7:**
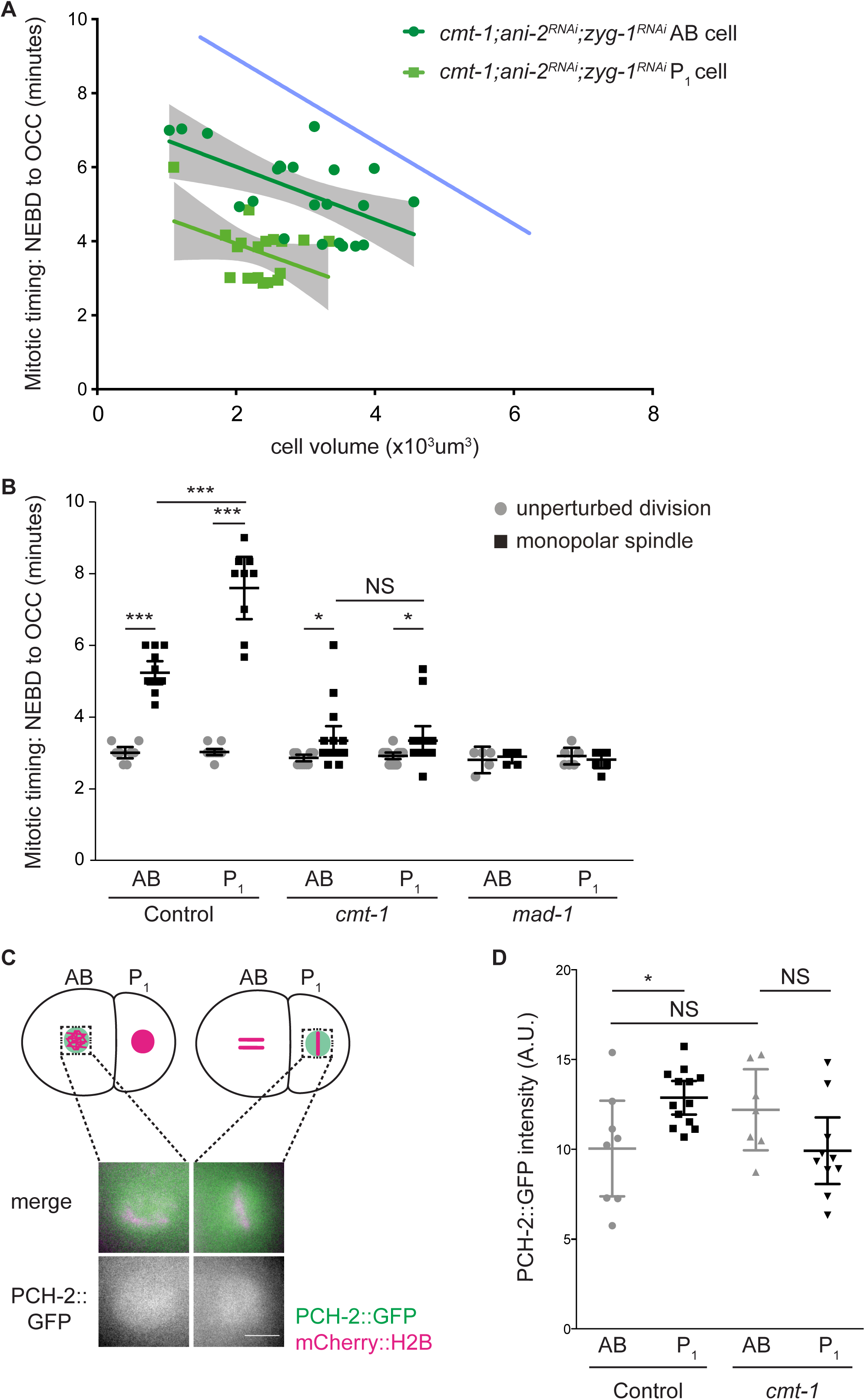
The stronger checkpoint in P_1_ cells depends on CMT-1. (A) Mitotic timing, as measured from nuclear envelope breakdown (NEBD) to the onset of cortical contractility (OCC), in AB and P_1_ cells plotted against cell volume in *cmt-1;ani-2*^*RNAi*^;*zyg-1*^*RNAi*^ embryos. Lines represent least-squares regression models with 95% confidence intervals (gray shaded areas) for each set of data. The opaque blue line represents the regression model of the control AB data from Figure 1C. Equations and p values indicating whether slopes are significantly non-zero for each model are: *cmt-1;ani-2*^*RNAi*^;*zyg-1*^*RNAi*^ AB (dark green): y=-0.713x+7.44 and p = 0.0050; *cmt-1;ani-2*^*RNAi*^;*zyg-1*^*RNAi*^ P_1_ (light green): y=-0.6767x+5.291 and p = 0.0452. (B) Mitotic timing of control, *cmt-1* and *mad-1* mutant embryos during unperturbed divisions or in the presence of monopolar spindles. (C) Cartoon and images of PCH-2::GFP localization around mitotic chromosomes in AB and P_1_ cells of *cmt-1* mutant embryos. Scale bar indicates 5 μm. (D) Quantification of PCH-2::GFP fluorescence in AB cells of control and *cmt-1* mutant embryos. (E) Quantification of PCH-2::GFP fluorescence in AB and P_1_ cells of *cmt-1* mutant embryos. All error bars are 95% confidence intervals. NS indicates not significant.

We also performed *zyg-1* RNAi on control and *cmt-1* mutant embryos and monitored mitotic timing in both AB and P_1_ cells. AB and P_1_ cells of control and *cmt-1* mutant embryos treated with control RNAi had similar mitotic timing. Unlike similar experiments in *pch-2* mutants (Figure 5C), we did not detect a statistically significant difference between cell cycle time in P_1_ cells between wildtype and *cmt-1* mutants embryos (Figure 7B), suggesting that *ani-2*^*RNAi*^;*zyg-1*^*RNAi*^ embryos might be more sensitive to subtle perturbations in cell cycle timing. In *zyg-1*^*RNAi*^ embryos, P_1_ cells exhibited a stronger checkpoint response than AB cells (Figure 7B). By contrast, both AB and P_1_ cells in *cmt-1*;*zyg-1*^*RNAi*^ mutant embryos exhibited similar spindle checkpoint delays (Figure 7B). Despite having spindle checkpoint responses that were less robust than that of control *zyg-1*^*RNAi*^ embryos, AB and P_1_ cells in *cmt-1* mutant embryos treated with *zyg-1* RNAi spent significantly longer in mitosis than *cmt-1* mutant embryos treated with control RNAi (Figure 7B), indicating that they activated a weaker spindle checkpoint response, similar to our published results (Nelson *et al*., 2015). More importantly, *cmt-1*;*zyg-1*^*RNAi*^ mutant embryos failed to generate a stronger checkpoint in P_1_ cells, consistent with *cmt-1;ani-2*^*RNAi*^;*zyg-1*^*RNAi*^ experiments (Figure 7A).

Aside from localizing PCH-2 to unattached kinetochores (Nelson *et al*., 2015), we wondered if CMT-1 was required for any other aspects of PCH-2 regulation. Therefore, we tested whether CMT-1 was necessary for PCH-2’s asymmetric enrichment in P_1_ cells. We quantified PCH-2::GFP fluorescence in prometaphase in the area around chromosomes in both *cmt-1* mutant AB and P_1_ cells (Figure 7C). First, we found that PCH-2::GFP fluorescence was slightly higher in AB cells in *cmt-1* mutants than control embryos (Figure 7D). We saw a similar result in our *par-1* RNAi experiments, although in both cases these increases were not statistically significant. However, unlike *par-1*^*RNAi*^ embryos (Gerhold *et al*., 2018), this increase in PCH-2::GFP was not accompanied by an increase in checkpoint strength (Figure 7B), consistent with our hypothesis that the weaker checkpoint in *cmt-1* AB cells is a consequence of PCH-2’s absence from unattached kinetochores (Nelson *et al*., 2015). Further, when we compared the quantification of PCH-2::GFP fluorescence in *cmt-1* mutant AB and P_1_ cells (Figure 7C), we did not detect a significant difference between the two cells (Figure 7D), unlike our experiment in control embryos (Figures 6B and C), indicating that CMT-1 contributes to the asymmetric enrichment of PCH-2 in P_1_ cells. Thus, CMT-1 promotes spindle checkpoint strength through two mechanisms: localizing PCH-2 to unattached kinetochores and ensuring its enrichment in germline precursor cells.

## DISCUSSION

The role of PCH-2, and its mammalian ortholog TRIP13, in the spindle checkpoint has been enigmatic. Originally identified as a checkpoint silencing factor (Eytan *et al*., 2014; Wang *et al*., 2014; Miniowitz-Shemtov *et al*., 2015; Ye *et al*., 2015; Brulotte *et al*., 2017; Alfieri *et al*., 2018), more recent evidence also indicates a role in promoting the checkpoint response (Nelson *et al*., 2015; Ma and Poon, 2016; Yost *et al*., 2017; Ma and Poon, 2018). It’s clear that the reliance on PCH-2/TRIP13 in checkpoint activation reflects the relative levels and availability of O-Mad2 (Ma and Poon, 2018). We show here that PCH-2 also controls checkpoint strength. Surprisingly, we can uncouple PCH-2’s requirement for checkpoint activation, which we detect in both AB and P_1_ cells of wildtype sized 2-cell embryos (Nelson *et al*., 2015 and Figure S4), from the requirement for spindle checkpoint strength, which we observe when we genetically manipulate cell size of 2-cell embryos by *ani-2* RNAi (Figures 1C and 5B). Based on this, we propose that PCH-2 regulates checkpoint strength not only by regulating O-Mad2 availability, but by doing so specifically at and near unattached kinetochores, providing an unanticipated mechanism to explain this phenomenon (Figure 8). This role in checkpoint strength appears to be particularly important in large cells, such as oocytes and cells in early embryos, as well as cells that give rise to immortal germ cells.

**Figure 8:**
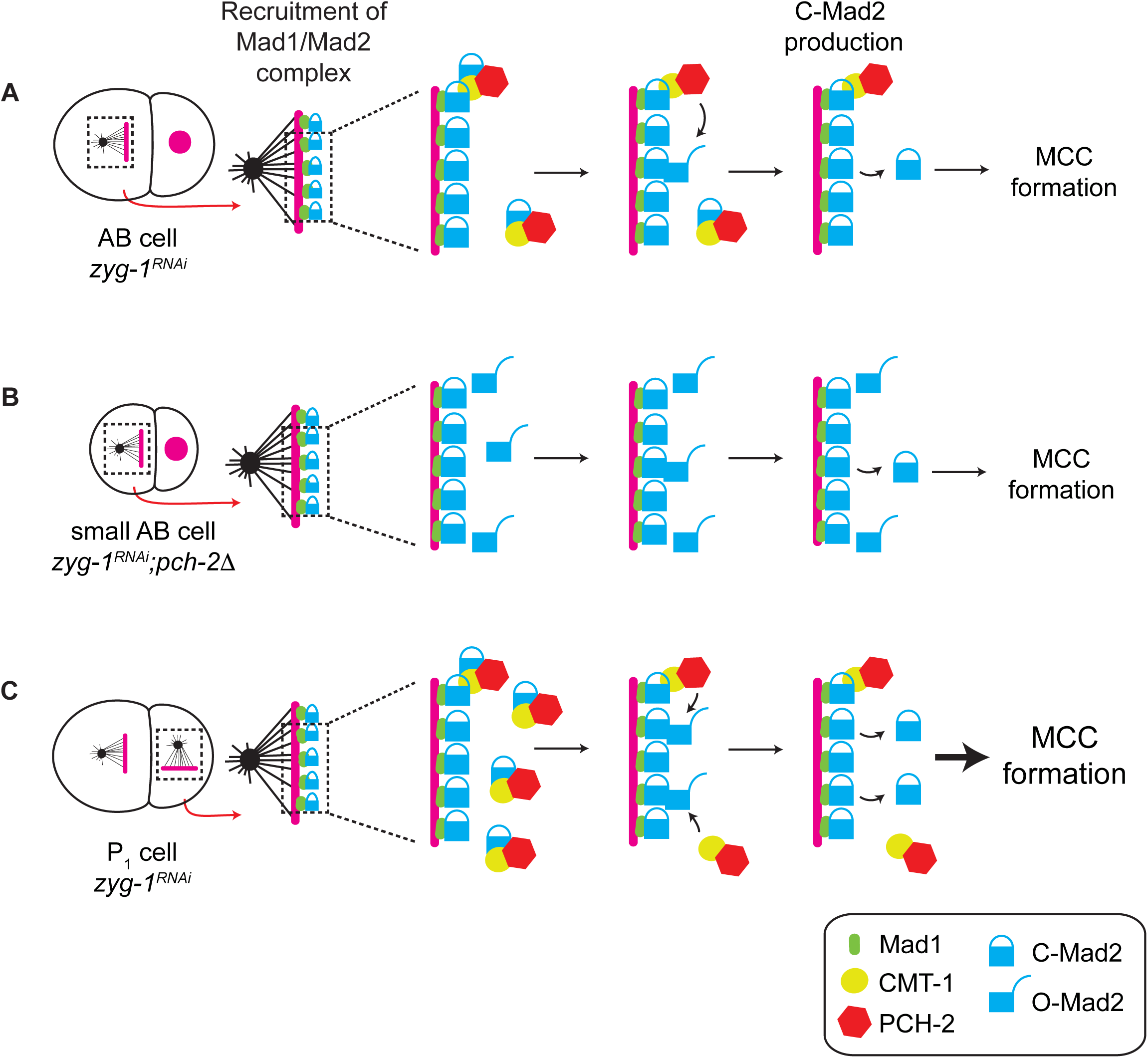
**Model** (A) A robust spindle checkpoint response in large cells requires the presence of PCH-2 at unattached kinetochores to increase the local concentration O-MAD-2 at and near unattached kinetochores. (B) Reducing cell volume of 2-cell embryos increases the concentration of O-Mad-2 at and near unattached kinetochores, allowing a checkpoint response in the absence of PCH-2. (C) The enrichment of PCH-2 around mitotic chromosomes in P_1_ cells results in a higher production of O-MAD-2, generating a stronger spindle checkpoint response in these cells.

Our model assumes that 2-cell embryos have a significant amount of O-Mad2 available, even when PCH-2 function is lost (Figure 8), unlike what is reported in mammalian cells (Ma and Poon, 2016). Given that this is a developmental system in which embryos have only undergone a single mitotic division before we perform our assays, we propose that O-Mad2 is not limiting in very early embryos, even in *pch-2* null mutants. In this way, *C. elegans* 2-cell embryos would be analogous to mammalian cells undergoing cell division soon after acute depletion of TRIP13 (Ma and Poon, 2018). Unfortunately, we were unable to directly probe O-Mad2 concentration or its availability at or near unattached kinetochores in small *ani-2*^*RNAi*^ embryos or germline precursor cells. However, we think that several pieces of data support our model (Figure 8). PCH-2’s characterized biochemical activity regulates the availability of O-Mad2 (Ye *et al*., 2015; Brulotte *et al*., 2017; Alfieri *et al*., 2018), making this the likely mechanism through which PCH-2 regulates checkpoint strength. PCH-2 at unattached kinetochores in AB and P_1_ cells (Nelson *et al*., 2015 and Figure 7B) and its enrichment around mitotic chromosomes in P_1_ cells (Figures 5A and 6C) correlates with a stronger checkpoint. The loss of PCH-2 or this enrichment produces similar checkpoint strength between AB and P_1_ cells (Gerhold et al., 2018 and Figures 1C, 5B, 6E, S7D, 7A and 7D). Indeed, the equalization of PCH-2::GFP between AB and P_1_ cells that we observe in *par-1*^*RNAi*^ and *par-6*^*RNAi*^ embryos (Figures 6E and S8D) is entirely consistent with the observation that in these mutants, AB cells more closely resemble P_1_ cells in spindle checkpoint strength (Gerhold et al., 2018). Finally, checkpoint strength depends on Mad2 dosage (Figure 2C), particularly in AB cells of 4-cell *pch-2* mutant embryos (Figure 4B).

Another prediction of our model is that overexpression of Mad2 should also make PCH-2 dispensable for spindle checkpoint activation. We’ve shown that subtle elevations of Mad2 protein levels introduce a cell cycle delay in AB cells of 4-cell embryos treated with nocodazole but not those treated with DMSO (Figure 4B), entirely consistent with our model. However, it’s not clear why we do not observe a similar effect in cells of the AB lineage of 8-cell or 16-cell embryos that overexpress Mad2 (Figure 4B). Unfortunately, more dramatic overexpression experiments are technically difficult in *C. elegans*. Further, it’s likely that strong overexpression of Mad2 in *C. elegans* embryos will delay normal mitosis, consistent with similar findings in mammalian cells (Marks *et al*., 2017) and budding yeast (Mariani *et al*., 2012). In this way, PCH-2’s function may provide a useful buffer: Since Mad2 protein levels may need to stay within a narrow range to allow normal mitotic timing, PCH-2’s localization at and near unattached kinetochores provide a mechanism to increase O-Mad2’s local concentration to promote effective and efficient signaling during checkpoint activation.

The requirement for PCH-2 in spindle checkpoint strength is also seen as AB cells normally decrease in volume during embryogenesis (Figure 4A), although not as dramatically as when we genetically manipulate cell size (Figure 1C). The inconsistency between our *ani-2* and embryogenesis experiments could be explained by a variety of factors. O-Mad2 may eventually become limiting in cells of the AB lineage with successive divisions after the 2-cell stage, resulting in a greater reliance on PCH-2 function. Moreover, it may also suggest that relative levels of O-Mad2 and C-Mad2 are more stringently regulated as embryonic development progresses and the multi-cellular embryo becomes more complex. This possibility is supported by our finding that PCH-2 regulates normal cell cycle timing in P_1_ cells, but not AB cells (Figure 5C), which implies that variations in O-Mad2/C-Mad2 ratios influence normal mitotic timing in cells with specific developmental fates. In addition, unlike the nuclei of 2-cell embryos treated with *ani-2*^*RNAi*^ (Figure S3), nuclear volume scales with cell volume during embryogenesis (Gerhold *et al*., 2018). Therefore, the concentration of Mad2 may not necessarily increase as cell size decreases in cells of the developing embryo, making direct comparisons between small cells obtained by *ani-2*^*RNAi*^ treatment and small cells resulting from normal embryogenesis challenging. Finally, recent reports have indicated that, during embryogenesis in other systems, cell volume may not be a major contributor to spindle checkpoint strength (Chenevert *et al*., 2019; Vazquez-Diez *et al*., 2019). Indeed, in *C. elegans*, when only AB cells are monitored during very early embryogenesis (the 2-8 cell stage), they exhibit very minor increases, if any, in checkpoint strength (Galli and Morgan, 2016; Gerhold *et al*., 2018 and Figure 4). This may suggest that cell fate is generally a more important determinant of spindle checkpoint strength during normal embryogenesis, potentially reconciling reports from a wide array of systems.

Our experiments identify CMT-1, the *C. elegans* ortholog of mammalian p31^comet^, as an important regulator of PCH-2 function and, as a result, checkpoint strength. In addition to its requirement in facilitating PCH-2’s ability to interact with its substrate, Mad2 (Miniowitz-Shemtov *et al*., 2015; Ye *et al*., 2015; Brulotte *et al*., 2017; Alfieri *et al*., 2018), CMT-1 localizes PCH-2 to unattached kinetochores (Nelson *et al*., 2015) and promotes PCH-2’s enrichment in P_1_ cells (Figure 7D). We propose that both of these roles contribute to checkpoint strength. In large AB cells, CMT-1 ensures PCH-2’s presence at unattached kinetochores, increasing the local concentration of O-Mad2, driving the production of soluble C-Mad2 and MCC and enforcing a robust checkpoint (Figure 8A). In P_1_ cells, the combination of PCH-2’s localization at kinetochores and its enrichment around chromosomes and near unattached kinetochores produces a checkpoint stronger than somatic cells (Figure 8C). It’s striking that, when CMT-1 is absent, AB cells, in which there is more PCH-2 (Figure 7D), and P_1_ cells, which are slightly smaller than AB cells, exhibit similar checkpoint strength (Figure 7B). This indicates that even these cells depend on PCH-2 to be present at unattached kinetochores to increase the local concentration of O-Mad2 and promote checkpoint strength.

P_1_ cells in both *pch-2;ani-2*^*RNAi*^;*zyg-1*^*RNAi*^ and *cmt-1;ani-2*^*RNAi*^;*zyg-1*^*RNAi*^ mutants show faster cell cycle timing than similarly sized AB cells of the same genotype (Figures 5B and 7A). However, only *pch-2* mutants significantly affect cell cycle timing in unperturbed P_1_ cells, (Figure 5C); P_1_ cells in *cmt-1* and *mad-1* mutants show accelerated cell cycle timing but this is not significantly faster than control (Figures 5C and 7B). Further, we don’t detect significant acceleration of the cell cycle in P_1_ cells of *pch-2;zyg-1*^*RNAi*^ or *mad-1;zyg-1*^*RNAi*^ mutant embryos (Figure S4 and 7B). Given the rapidity of cell cycles in these early embryos, it’s possible that *ani-2*^*RNAi*^;*zyg-1*^*RNAi*^ experiments provide greater sensitivity to observe subtle accelerations in cell cycle timing and that some subset of spindle checkpoint components, including PCH-2, CMT-1, MAD-1 and MAD-2 regulate normal cell cycle timing in germline precursor cells, similar to the role of MAD-1 and MAD-2 in germline mitotic nuclei (Gerhold *et al*., 2015). Unfortunately, we cannot test this with MAD-1 or MAD-2 since *mad-1* and *mad-2* mutants abolish the spindle checkpoint response in *ani-2*^*RNAi*^;*zyg-1*^*RNAi*^ embryos (Gerhold *et al*., 2015) and Figure 1C). An alternative hypothesis that we do not favor is that only PCH-2 regulates cell cycle timing in P_1_ cells, in a mechanism independent of other spindle checkpoint proteins.

Evolutionary analysis across phyla have revealed a close co-evolutionary relationship between PCH-2 and its orthologs and HORMA domain containing proteins, including CMT-1 and Mad2 (Vleugel *et al*., 2012; van Hooff *et al*., 2017). However, some organisms that rely on the templated conversion of O-Mad2 to C-Mad2 to assemble the MCC, such as budding and fission yeasts (Nezi *et al*., 2006; Chao *et al*., 2012) either don’t express their PCH-2 ortholog during mitosis (budding yeast) (San-Segundo and Roeder, 1999) or don’t have a PCH-2 ortholog in their genome (fission yeast) (Wu and Burgess, 2006). This is potentially explained by cell volume: Both budding and fission yeasts are two orders of magnitude smaller than mammalian cells and *C. elegans* embryos. They also undergo closed mitosis, in which the nuclear envelope does not break down, providing an additional opportunity to concentrate factors required for mitosis. We propose that recruiting O-Mad2 to unattached kinetochores may not present as great a challenge in these significantly smaller cells, making a factor required to increase the local concentration of O-Mad2 at unattached kinetochores unnecessary.

An obvious question our experiments raise is how PCH-2 is enriched in P_1_ cells. Germline precursor cells are transcriptionally silent until gastrulation (Seydoux *et al*., 1996) and sequencing of mRNA in early embryos shows that both CMT-1 and PCH-2 mRNA are not enriched in germline precursor cells (Tintori *et al*., 2016), indicating that enrichment of PCH-2 is likely to occur at the level of protein regulation. Understanding this regulation, its control by developmental events and its effect on the relative levels of O-Mad2 and C-Mad2 in different cell types promises to be an exciting area of investigation.

## MATERIALS AND METHODS

### Worms strains

The *C. elegans* Bristol N2 (Brenner, 1974) was used as the wild-type strain. Most strains were maintained at 20°C, except for *zyg-1(or297)* strains, which were maintained at 15°C. See Table S1 for the list of all the strains used in this study.

### Immunostaining

Immunostaining was performed on adult worms 48h after L4, as described in (Bhalla and Dernburg, 2005). The antibodies used were rabbit anti-MAD-2 (1/500; (Essex *et al*., 2009) and mouse anti-MAb414 (1/400; (Davis and Blobel, 1986). Secondary antibodies were Alexa Fluor 488 anti-rabbit (Invitrogen) and Cy3 anti-mouse (Jackson ImmunoResearch Laboratories, Inc.) diluted at 1:500. Antibody against MAD-2 was a gift from A. Desai (Ludwig Institute/University of California, San Diego, La Jolla, CA).

Images were acquired on a DeltaVision Personal DV microscope (GE Healthcare) equipped with a 100× NA 1.40 oil-immersion objective (Olympus), a short ARC xenon lamp (GE Healthcare) and using a CoolSNAP charge-coupled camera (Roper Scientific). Z-stacks were collected at 0.2 µm Z-spacing and processed by constrained, iterative deconvolution. Imaging, image scaling, and analysis were performed using functions in the softWoRx software package (GE Healthcare). Projections were calculated by a maximum intensity algorithm. Composite images were assembled and some false coloring was performed with Fiji.

### Live imaging of 2-cell embryos

For live imaging of 2-cell embryos, worms were dissected on glass coverslips in egg buffer and then mounted on 2% agar pads. Images were acquired every 1 minute or 20 seconds on a DeltaVision Personal DV microscope as described in the previous section; except that the distance between two planes was 2 µm. Mitotic timing was measured from NEBD to OCC as described in (Nelson *et al*., 2015). Cell volumes were measured as described in (Galli and Morgan, 2016). To measure the nuclear area, a sum projection of the embryo was generated 1 minute before chromosomes began to condense and the area was measured with Fiji (Figure S3A).

### Live imaging of embryogenesis

After treatment with *perm-1*^*RNAi*^ (see below), worms were dissected onto a coverslip with egg salt buffer (118 mM NaCl, 48 mM KCl) supplemented with 10 mM PIPES pH 7.3, 1 mM ATP and 10 mM sucrose. Embryos and adult carcasses were transferred into a well of an 8-well plate (ibidi 1 μ-Slide 8 Well Glass bottom) that had been freshly coated with 0.1% Poly-L-Lysine solution (Sigma P8920) and extensively washed. Time-lapse videos were acquired with a Solamere spinning disk confocal system piloted by μManager software (Edelstein *et al*., 2014) and equipped with a Yokogawa CSUX-1 scan head, a Nikon (Garden City, NY) TE2000-E inverted stand, a Hamamatsu ImageEM ×2 camera, LX/MAS 489 nm and LS/MAS 561 nm laser, and Plan Apo ×60/1.4 numerical aperture oil objective. Acquisition times per frame were 50 ms using 5% of the lasers power for both channels, and images were obtained as stacks of planes at 2 μm intervals taken every 1 minute. Nocodazole was added from a 5X stock to a final concentration of 50 μM after the first time point. Mitotic timing was measured from NEBD to DECON as described in (Essex *et al*., 2009).

To image embryogenesis in *zyg-1(or297)* mutants, images were generated under the same conditions as previously described for the live imaging of 2-cell embryos with a few modifications: Images were acquired every 20 seconds on a DeltaVision Personal DV microscope in a room heated to 26°C. Mitotic timing was measured from NEBD to DECON as described in (Essex *et al*., 2009).

### Quantification of fluorescence intensity

To quantify GFP::MAD-2 and PCH-2::GFP levels, images were generated under the same conditions as previously described for the live imaging of 2-cell embryos with a few modifications: only the area defined by the GFP cloud and mitotic chromosomes was imaged, the interval between the four planes was 1 μm and images were collected every 20 seconds. Quantification of fluorescence at kinetochores was performed in Fiji as described in (Moyle *et al*., 2014; Nelson *et al*., 2015). Briefly, maximum intensity projections of both mCh::H2B and GFP fusion proteins were made after the pseudo-metaphase plate was generated. The image was rotated so the metaphase plate was vertical, channels were split, and the maximum GFP pixel was identified using the process function within a box on the unattached side of the metaphase plate. In the same x-plane, the maximum mCh::H2B pixel was found. The width was changed to 12 pixels and the maximum GFP signal intensity was recorded in this 12 pixel window centered at the mCherry maxima. The background GFP signal was calculated by taking the average GFP intensity of a 4 pixel box in the same x-plane, 8 pixels away from the maximum mCherry on the opposite side of the pseudo-metaphase plate to the maximum GFP (i.e. the attached side). This background GFP was then subtracted from the maximum to measure the kinetochore bound GFP fusion intensity. Fluorescence around mitotic chromosomes was quantified as described in (Galli and Morgan, 2016). Sum intensity projections were generated and fluorescence in the area around mitotic chromosomes was measured in Fiji. Background fluorescence was measured in a 30 pixel band around this “cloud” and subtracted from the initial fluorescence intensity to determine the final fluorescence value. In some of our movies, identifying a clear metaphase plate was more difficult in AB than P_1_ cells. Therefore, to ensure that we were quantifying PCH-2::GFP fluorescence around mitotic chromosomes at the same stage in mitosis in these two cell types, PCH-2::GFP was quantified in frames that were normalized relative to NEBD and mitotic exit. To measure the cell volume, one Z-stack of the entire cell was taken at NEBD at 2 µm Z-spacing.

### Feeding RNA interference (RNAi)

*C. elegans* strains were fed HT115 bacteria expressing the desired dsRNA after IPTG induction. Bacterial strains containing RNAi vectors were cultured overnight at 37°C, centrifuged, and the pellet was resuspended in 1/50 of the original volume. 100 µl of concentrated culture was spotted onto a nematode growth medium (NGM) plate containing 1 mM IPTG and 50 µg/μl of kanamycin or carbenicillin and the plate was incubated overnight at 37°C.

For *ani-2* RNAi, gravid adults were bleached onto the RNAi plate and their progeny was allowed to develop at 20°C during 2.5 days. Then, L4s were transferred to a fresh plate containing OP50 or *zyg-1* RNAi bacteria.

For *zyg-1* RNAi, L4s were transferred (from an OP50 or *ani-2* RNAi plate) onto a *zyg-1* RNAi plate and cultured 1.5 days at 20°C.

For *perm-1* RNAi, young adults (8h post L4) were incubated onto *perm-1* RNAi plates for 16-20 hours at 15°C.

For *par-1, par-6* and *gpr-1/2* RNAi, gravid adults were bleached onto control RNAi (L4440) plates and their progeny were allowed to develop at 15°C for 3 days. L4s were then transferred onto *par-1, par-6, gpr-1/2* RNAi or control RNAi plates and incubated at 15°C for 3 days. For *gpr-1/2* RNAi, “small” AB cells were identified by whether their area was at least one standard deviation lower than the average of control AB cells and “large” P_1_ cells were identified by whether their area was at least one standard deviation higher than the average of control P_1_ cells.

### Statistical Analysis

All statistical analysis was performed using GrapPad Prism version 6 for Macintosh, including linear regression analysis and assessing significance of this data (Figures 1C, 2B, 3C, 5A, 5B, 7A, S1C, and S1D). For comparing two means, significance was assessed by performing two-tailed t-tests (Figures 6C, S5B, S5D, S6B, S6C, S6D, and S7A). In graphs in which multiple means were tested, we performed ANOVA analysis with the Sidak post-hoc test (Figures 1D, 5C, 6A, 6E, 7B, 7D, S4A, S8A, S8B, and s8D). In all graphs, a * indicates a p value < 0.05, ** indicates a p value < 0.01 and *** a p value < 0.0001.

## ACKNOWLEDGEMENTS

We would like to thank Arshad Desai, Karen Oegema, Susan Strome and David Morgan for valuable strains and reagents. This work was supported by the NIH (grant numbers T32GM008646 [C.R.N. and A.R.] and R01GM097144 [N.B.]). Some strains were provided by the CGC, which is funded by NIH Office of Research Infrastructure Programs (P40 OD010440).

## CONTRIBUTIONS

Conceptualization and Methodology, L.D., A.E.R., C.R.N. and N.B.; Investigation, L.D., A.E.R., and C.R.N.; Writing - Original Draft, L.D. and N.B.; Writing - Review & Editing, L.D., A.E.R., C.R.N. and N.B.; Supervision and Funding Acquisition, N.B.

---

**Figure S1:**
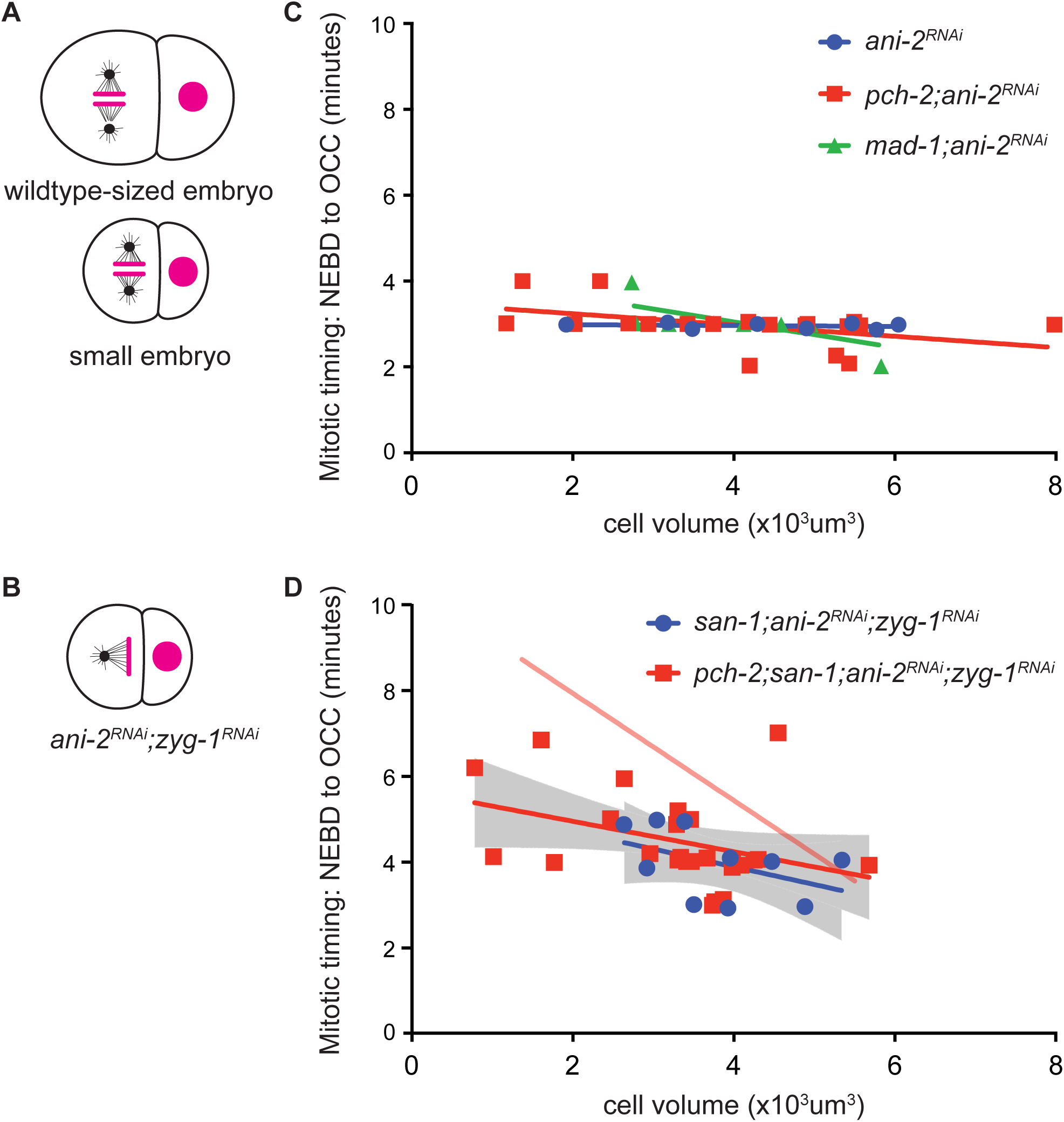
Related to Figure 1. The mitotic delay observed in *pch-2;ani-2*^*RNAi*^;*zyg-1*^*RNAi*^ small cells is a spindle checkpoint response. Cartoon of control and *ani-2*^*RNAi*^ 2-cell embryos (A) and an *ani-2*^*RNAi*^;*zyg-1*^*RNAi*^ 2-cell embryo (B). Mitotic timing in AB cells plotted against cell volume, during unperturbed mitosis (C) and in the presence of monopolar spindles. (D). Lines represent least-squares regression models for each set of data. For (C) equations and p values indicating whether slopes are significantly non-zero for each model are: *ani-2*^*RNAi*^ (blue): y=-9.744×10^−3^x+3.001 and p = 0.6073; *pch-2;ani-2*^*RNAi*^ (red): y=-0.1315x+3.503 and p = 0.0483; *mad-1;ani-2*^*RNAi*^ (green): y=-0.2944x+4.247 and p = 0.0251. In (D), 95% confidence intervals are indicated by gray shaded areas and equations and p values indicating whether slopes are significantly non-zero for each model are: *san-1;ani-2*^*RNAi*^;*zyg-1*^*RNAi*^ P_1_ (blue): y=-0.4136x+5.535 and p = 0.1885; *san-1;pch-2;ani-2*^*RNAi*^;*zyg-1*^*RNAi*^ P_1_ (red): y=-0.3541x+5.647 and p = 0.0643. The regression model of *pch-2;ani-2*^*RNAi*^;*zyg-1*^*RNAi*^ embryos from Figure 1C is indicated by the opaque red line in (D) for comparison.

**Figure S2:**
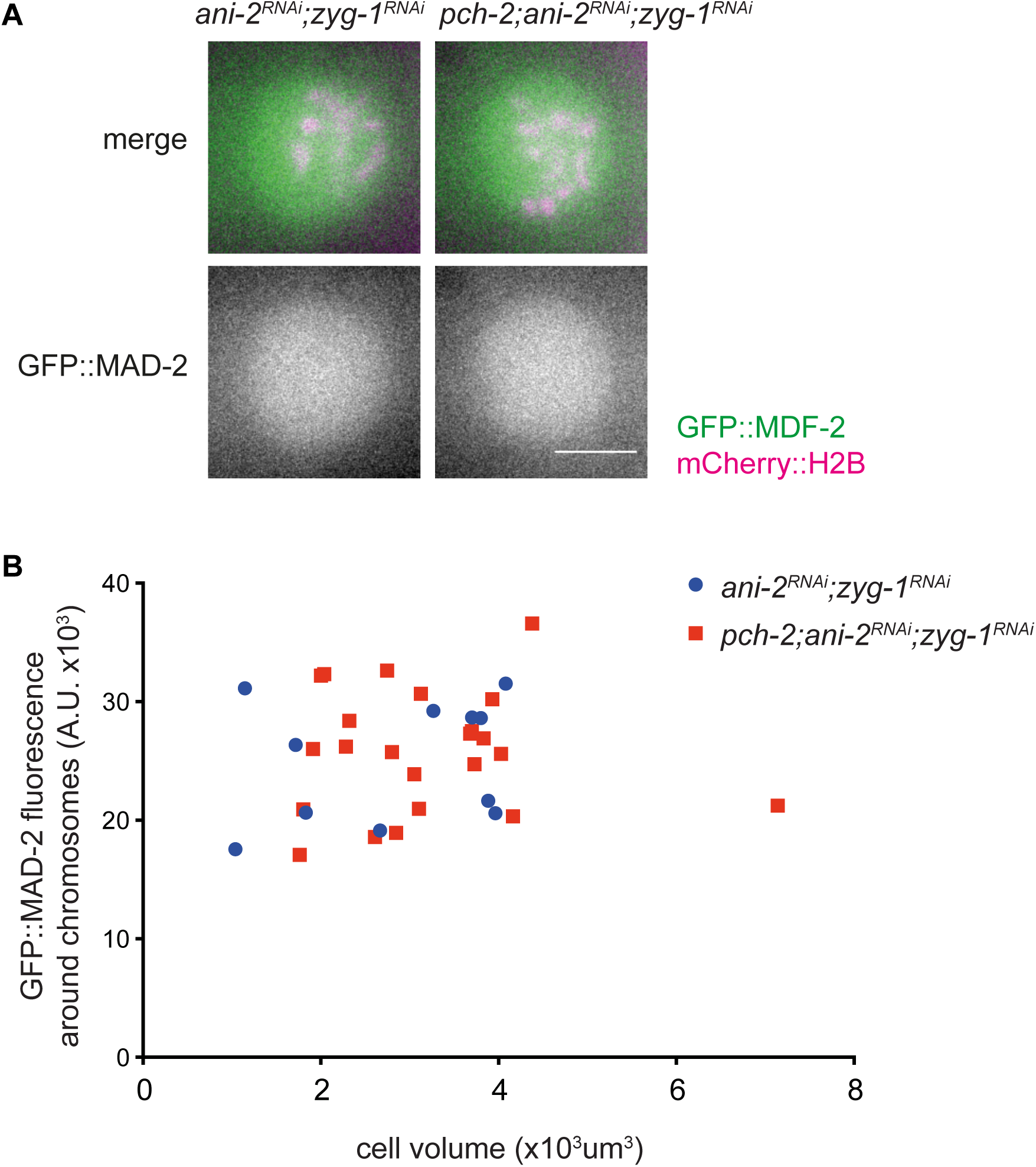
Related to Figure 2: There is no difference in GFP::MAD-2 fluorescence around mitotic chromosomes in *ani-2*^*RNAi*^;*zyg-1*^*RNAi*^ embryos and *pch-2;ani-2*^*RNAi*^;*zyg-1*^*RNAi*^ AB cells. (A) Images of MAD-2::GFP in AB cells of control *ani-2*^*RNAi*^;*zyg-1*^*RNAi*^ embryos or *pch-2;ani-2*^*RNAi*^;*zyg-1*^*RNAi*^ embryos after NEBD. Scale bar indicates 5 μm. (B) Quantification of GFP::MAD-2 fluorescence around mitotic chromosomes in AB cells of control and *pch-2* embryos plotted against cell volume.

**Figure S3:**
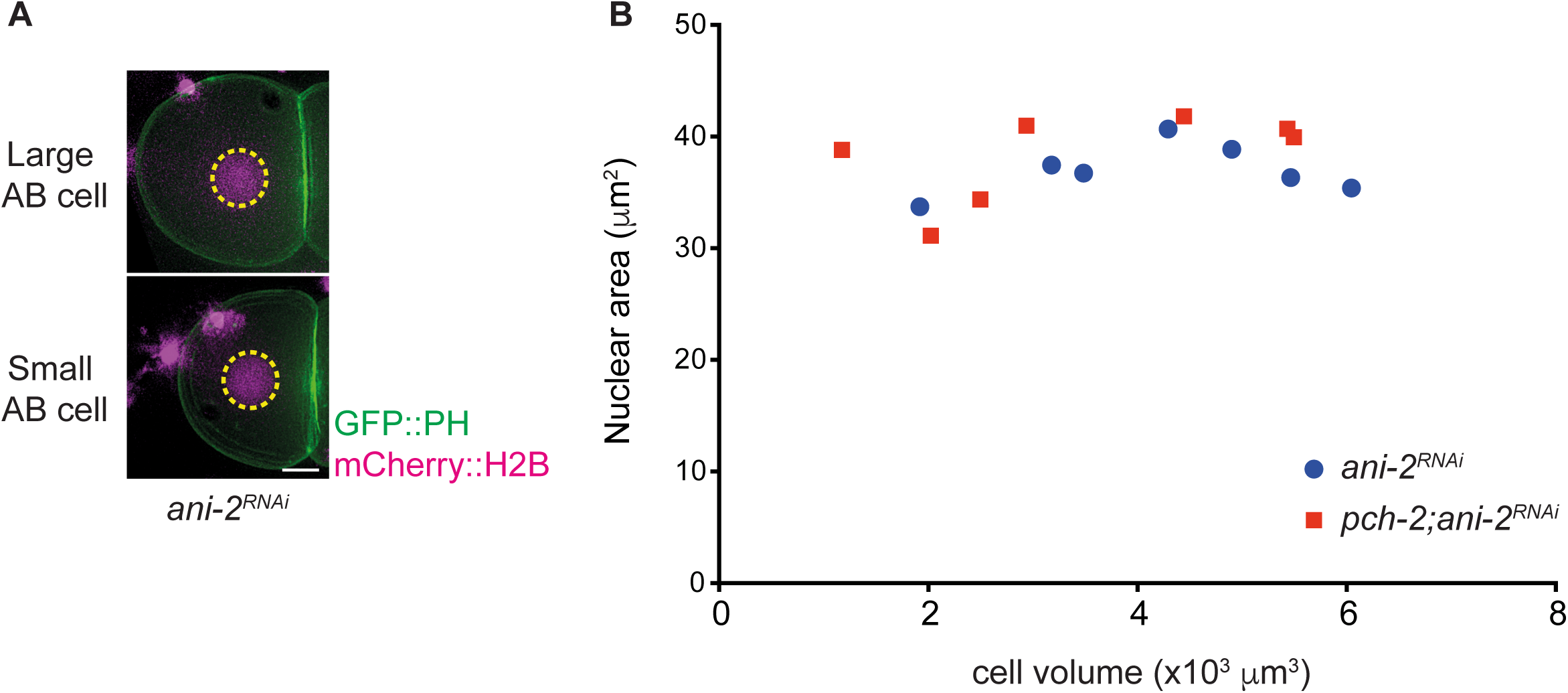
Related to Figure 3. Nuclear volume does not scale with cell volume in *ani-2*^*RNAi*^ 2-cell embryos. (A) Images of a large (top) and small (bottom) AB cell of *ani-2*^*RNAi*^ embryos. The nuclear area is indicated with a dashed yellow line. Scale bar indicates 5 μm. (B) Nuclear area plotted against cell volume.

**Figure S4:**
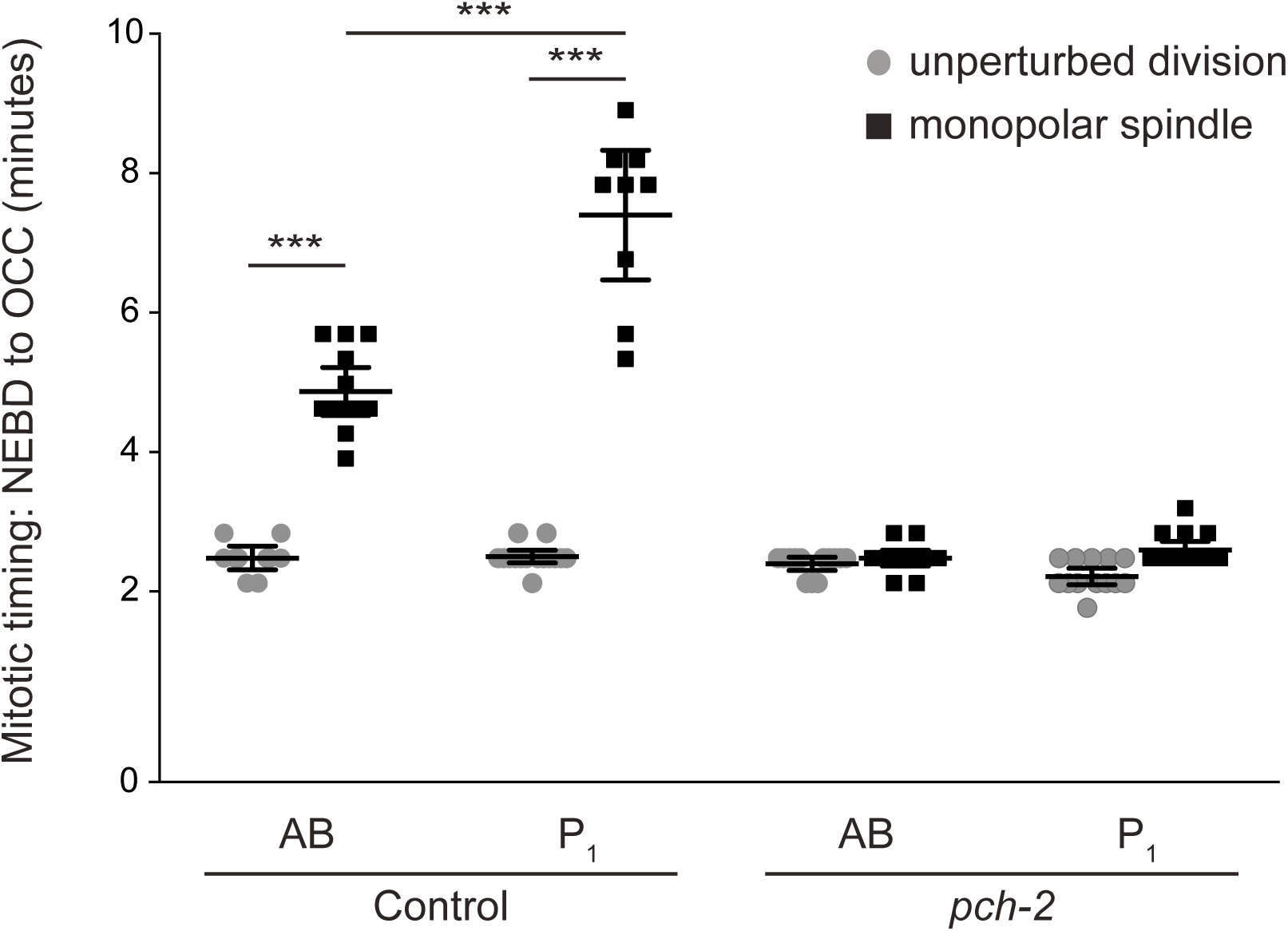
Related to Figure 5. PCH-2 is required for the spindle checkpoint in the germline lineage. (A) Mitotic timing of control and *pch-2* mutant embryos during unperturbed divisions or in the presence of monopolar spindles. Data for control embryos is the same as Figure 7B. Error bars indicate 95% confidence intervals.

**Figure S5:**
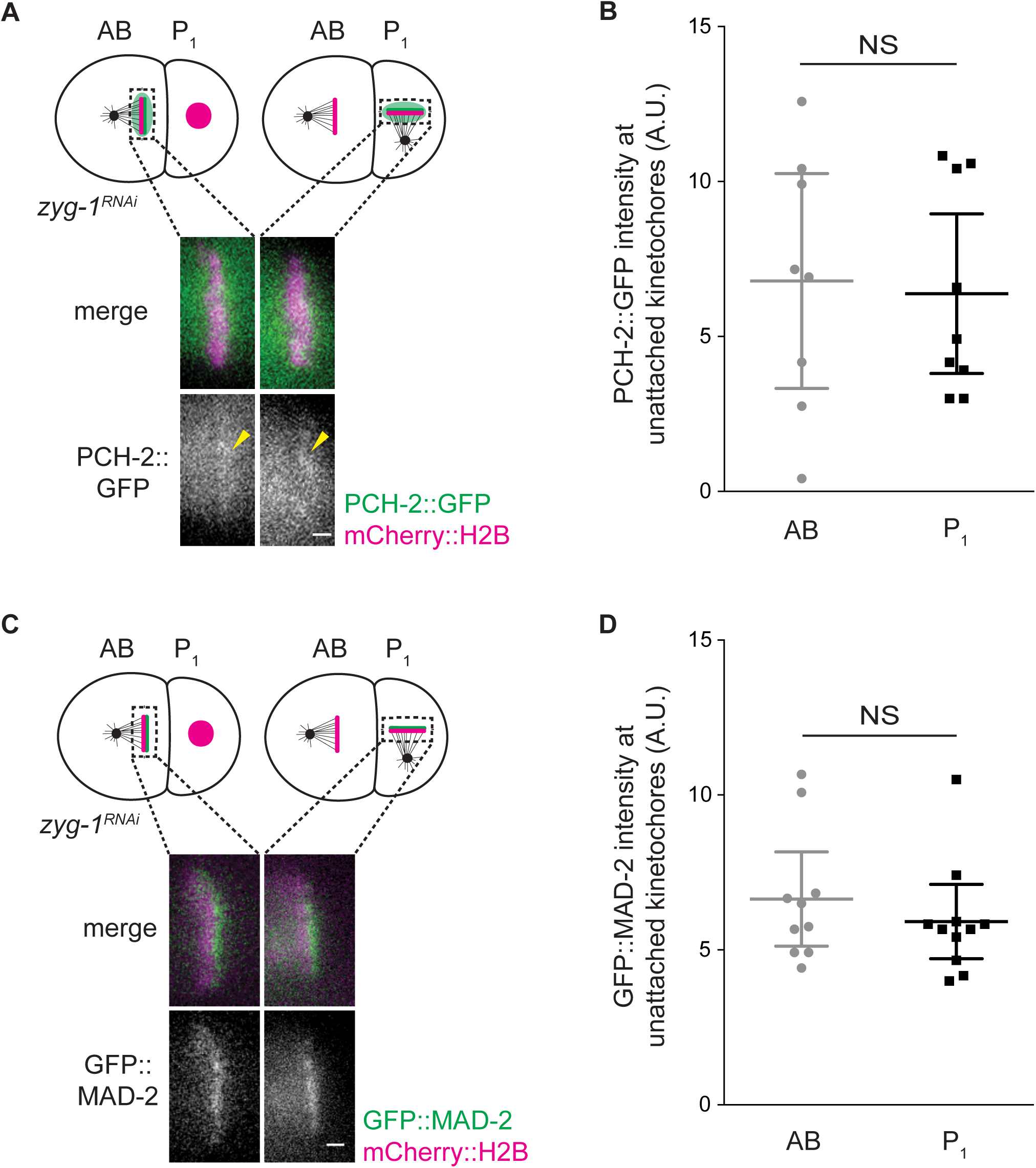
Related to Figure 6. There is no difference in the amount of PCH-2::GFP or GFP::MAD-2 recruited to unattached kinetochores in AB and P_1_ cells. (A) Cartoon and images of PCH-2::GFP recruitment to unattached kinetochores in AB and P_1_ cells of 2-cell embryos. Scale bar indicates 1 μm. (B) Quantification of PCH-2::GFP recruitment at unattached kinetochores in AB and P_1_ cells. (C) Cartoon and images of GFP::MAD-2 recruitment to unattached kinetochores in AB and P_1_ cells of 2-cell embryos. Scale bar indicates 1 μm. (D) Quantification of GFP::MAD-2 fluorescence at unattached kinetochores in AB and P_1_ cells. All error bars are 95% confidence intervals. NS indicates not significant.

**Figure S6:**
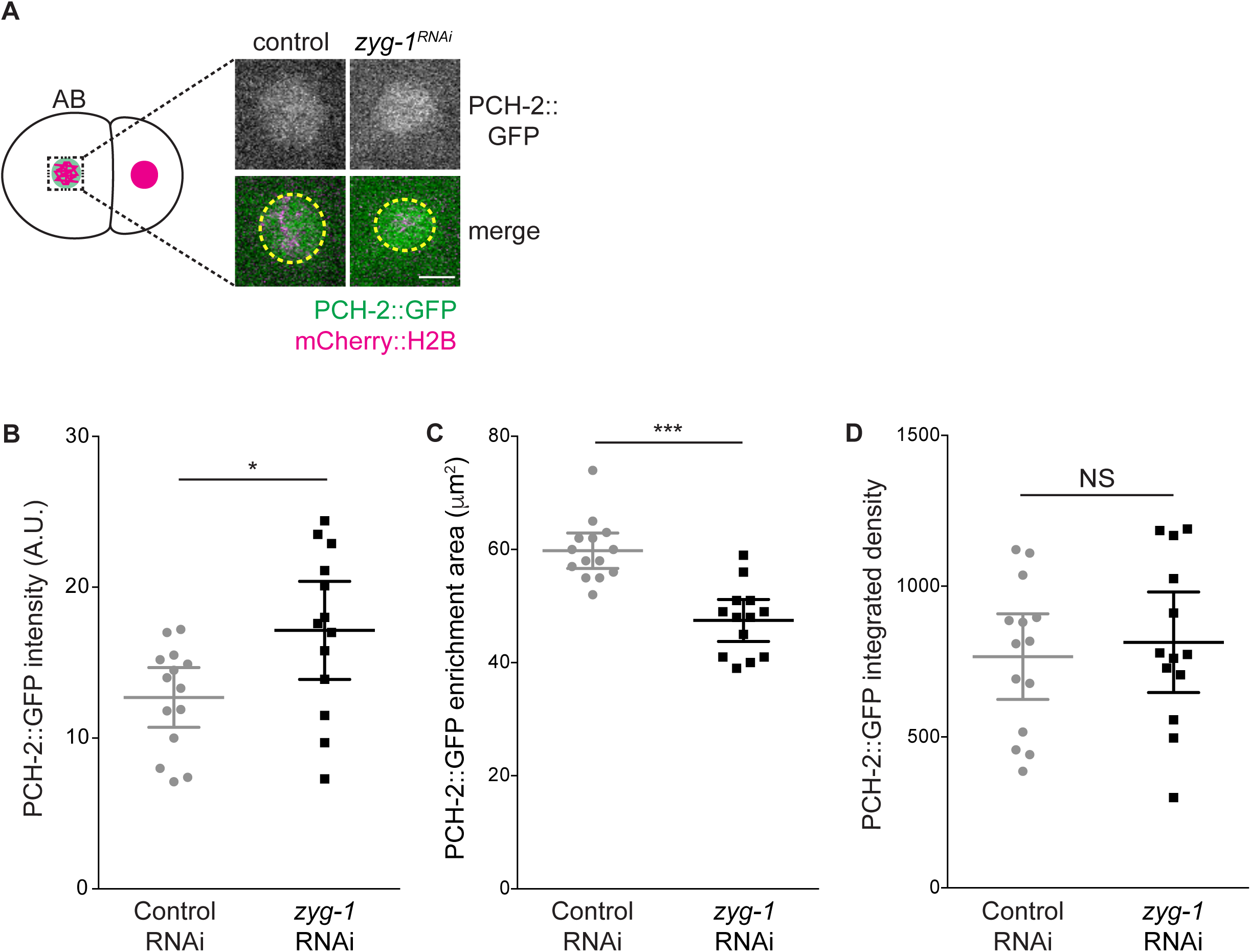
Related to Figure 6. There is no difference in the amount of PCH-2::GFP around mitotic chromosomes in AB cells with bipolar or monopolar spindles. (A) Cartoon and images of PCH-2::GFP localization around mitotic chromosomes in AB cells of 2-cell embryos. Scale bar indicates 5 μm. Yellow dashed circle indicates area of PCH-2::GFP fluorescence. (B) Quantification of PCH-2::GFP fluorescence in AB cells with bipolar spindles (control) or monopolar spindles (*zyg-1*^*RNAi*^). (C) Quantification of area of PCH-2::GFP fluorescence in AB cells with bipolar spindles or monopolar spindles. (D) Quantification of integrated density of PCH-2::GFP fluorescence in AB cells with bipolar spindles or monopolar spindles. All error bars are 95% confidence intervals. NS indicates not significant.

**Figure S7:**
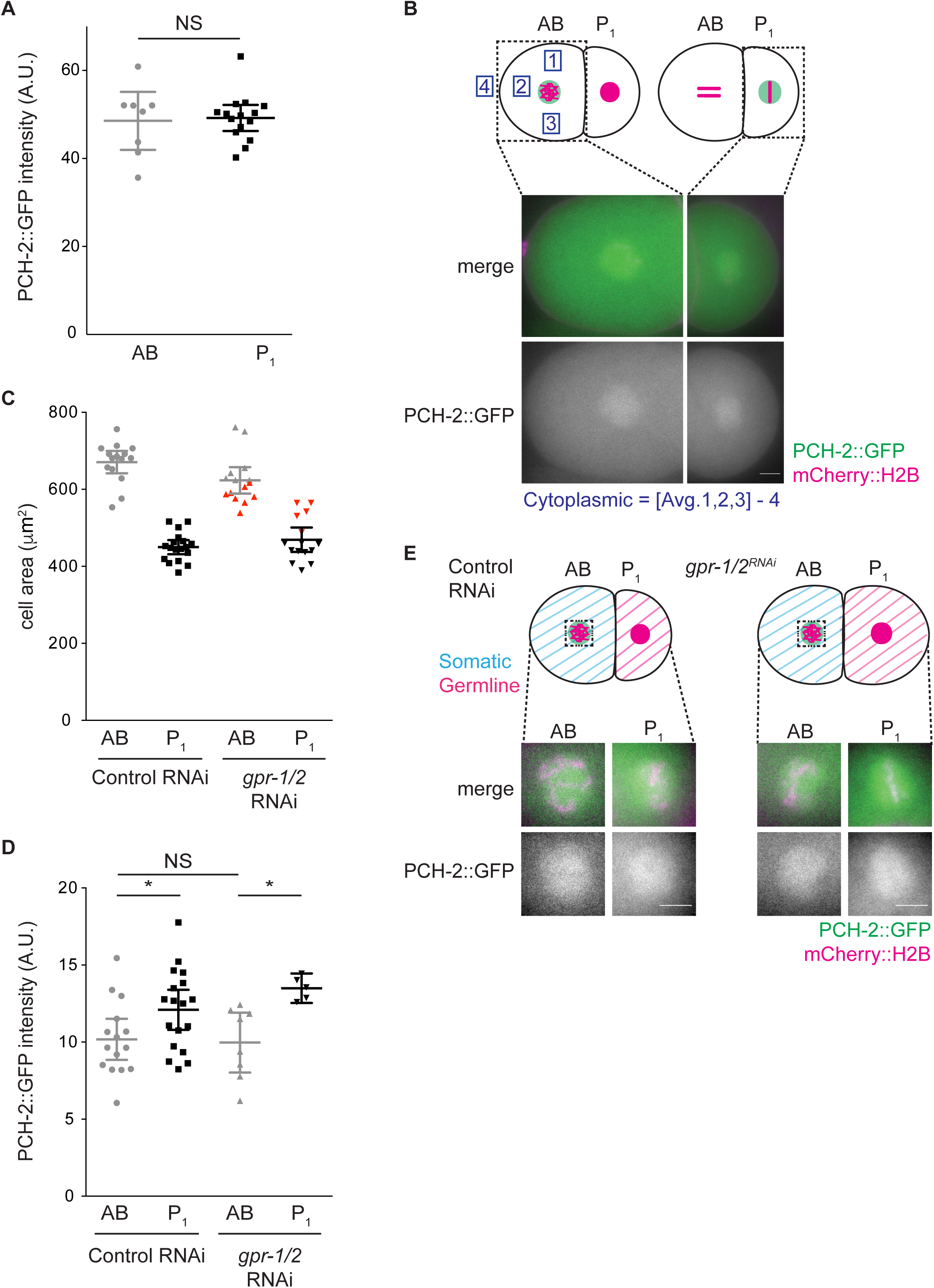
Related to Figure 6. PCH-2’s enrichment in P1 cells is around chromosomes and does not depend on GPR-1/2. (A) Quantification of PCH-2::GFP fluorescence in the cytoplasm of AB and P_1_ cells. (B) Images of AB (left) and P_1_ (right) cells after NEBD. Scale bar indicate 5 μm. (C) Quantification of cell area in AB and P_1_ cells of control RNAi and *gpr-1/2*^*RNAi*^ 2-cell embryos. Red symbols indicate cells in which PCH-2::GFP fluorescence was quantified in (D). (D) Quantification of PCH-2::GFP fluorescence in AB and P_1_ cells of control RNAi and *gpr-1/2*^*RNAi*^ embryos. (E) Cartoon and images of PCH-2::GFP localization around mitotic chromosomes in AB and P_1_ cells of control RNAi and *gpr-1/2*^*RNAi*^ 2-cell embryos. Scale bars indicate 5 μm. Error bars are 95% confidence intervals. NS indicates not significant.

**Figure S8:**
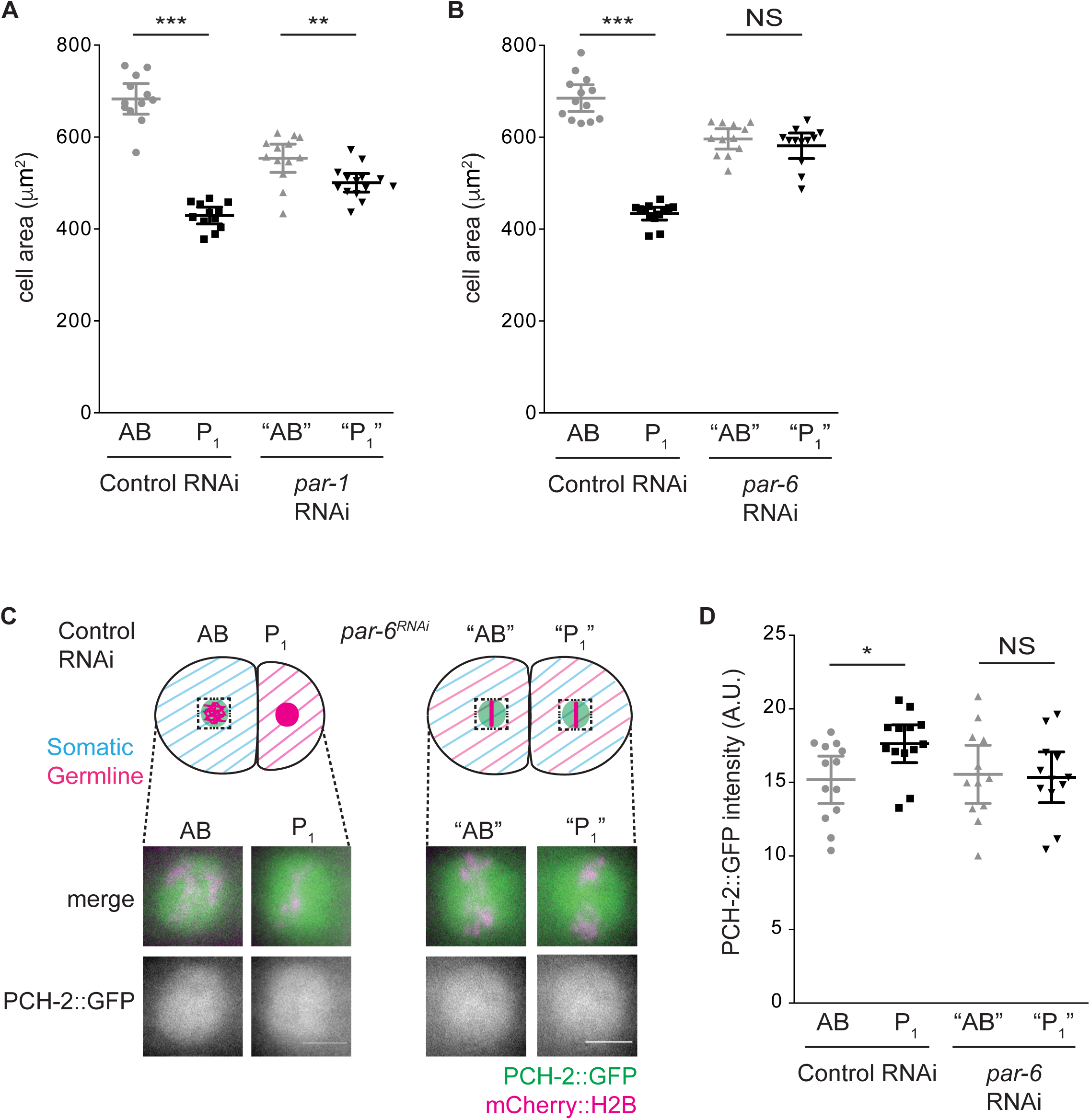
Related to Figure 6. PCH-2’s enrichment around mitotic chromosomes in P_1_ cells depends on PAR-6. (A and B) Quantification of cell area in AB and P_1_ cells of control RNAi, *par-1*^*RNAi*^ and *par-6*^*RNAi*^ 2-cell embryos. (C) Cartoon and images of PCH-2::GFP localization around mitotic chromosomes in AB and P_1_ cells of control RNAi and *par-6*^*RNAi*^ 2-cell embryos. Scale bars indicate 5 μm. (D) Quantification of PCH-2::GFP fluorescence in AB and P_1_ cells of control RNAi and *par-6*^*RNAi*^ embryos. Error bars are 95% confidence intervals. NS indicates not significant.

## Movie legends

**Video 1**

Mitosis in the AB cell of a wild-type sized control 2-cell embryo with monopolar spindles expressing GFH::PH and mCherry::H2B for visualization of the plasma membrane and the chromosomes, respectively (strain OD95). The timer starts at NEBD.

**Video 2**

Mitosis in the AB cell of a small control 2-cell embryo with monopolar spindles expressing GFH::PH and mCherry::H2B for visualization of the plasma membrane and the chromosomes, respectively (strain OD95). The timer starts at NEBD.

**Video 3**

Mitosis in the AB cell of a wild-type sized *pch-2(tm1458)* 2-cell embryo with monopolar spindles expressing GFH::PH and mCherry::H2B for visualization of the plasma membrane and the chromosomes, respectively (strain BHL575). The timer starts at NEBD.

**Video 4**

Mitosis in the AB cell of a small *pch-2(tm1458*) 2-cell embryo with monopolar spindles expressing GFH::PH and mCherry::H2B for visualization of the plasma membrane and the chromosomes, respectively (strain BHL575). The timer starts at NEBD.

**Video 5**

Mitosis in the P_1_ cell of a wild-type sized control 2-cell embryo with monopolar spindles expressing GFH::PH and mCherry::H2B for visualization of the plasma membrane and the chromosomes, respectively (strain OD95). The timer starts at NEBD.

**Video 6**

Mitosis in the P_1_ cell of a small control 2-cell embryo with monopolar spindles expressing GFH::PH and mCherry::H2B for visualization of the plasma membrane and the chromosomes, respectively (strain OD95). The timer starts at NEBD.

**Video 7**

Mitosis in the P_1_ cell of a wild-type sized *pch-2(tm1458)* 2-cell embryo with monopolar spindles expressing GFH::PH and mCherry::H2B for visualization of the plasma membrane and the chromosomes, respectively (strain BHL575). The timer starts at NEBD.

**Video 8**

Mitosis in the P_1_ cell of a small *pch-2(tm1458)* 2-cell embryo with monopolar spindles expressing GFH::PH and mCherry::H2B for visualization of the plasma membrane and the chromosomes, respectively (strain BHL575). The timer starts at NEBD.

**Video 9**

Mitosis in the AB cell of a wild-type sized *cmt-1(ok2879*) 2-cell embryo with monopolar spindles expressing GFH::PH and mCherry::H2B for visualization of the plasma membrane and the chromosomes, respectively (strain BHL608). The timer starts at NEBD.

**Video 10**

Mitosis in the AB cell of a small *cmt-1(ok2879*) 2-cell embryo with monopolar spindles expressing GFH::PH and mCherry::H2B for visualization of the plasma membrane and the chromosomes, respectively (strain BHL608). The timer starts at NEBD.

**Video 11**

Mitosis in the P_1_ cell of a wild-type sized *cmt-1(ok2879*) 2-cell embryo with monopolar spindles expressing GFH::PH and mCherry::H2B for visualization of the plasma membrane and the chromosomes, respectively (strain BHL608). The timer starts at NEBD.

**Video 12**

Mitosis in the P_1_ cell of a small *cmt-1(ok2879*) 2-cell embryo with monopolar spindles expressing GFH::PH and mCherry::H2B for visualization of the plasma membrane and the chromosomes, respectively (strain BHL608). The timer starts at NEBD.

**Table S1:**
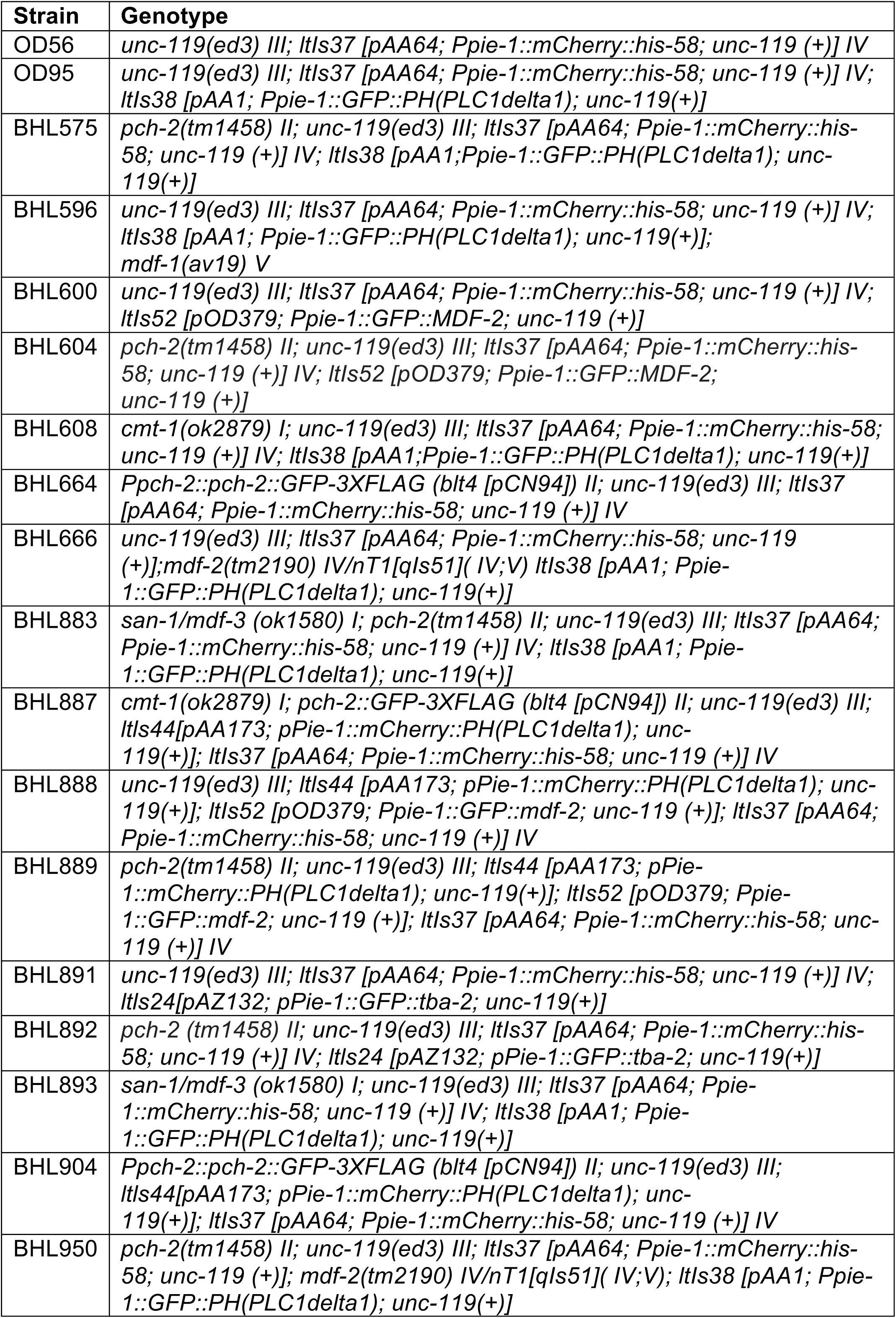
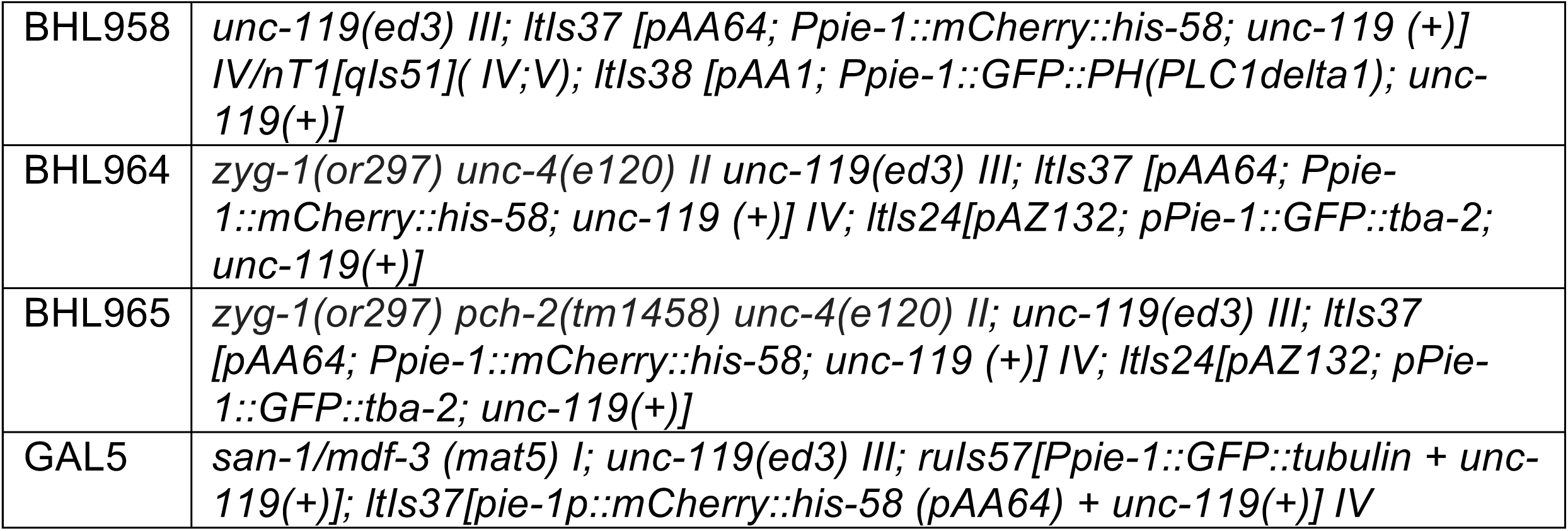
Strains used in this study.

| Strain | Genotype |
| --- | --- |
| OD56 | *unc-119(ed3) III; ItIs37 [pAA64; Ppie-1::mCherry::his-58; unc-119 (+)] IV* |
| OD95 | *unc-119(ed3) III; ItIs37 [pAA64; Ppie-1::mCherry::his-58; unc-119 (+)] IV; ItIs38 [pAA1; Ppie-1::GFP::PH(PLC1delta1); unc-119(+)]* |
| BHL575 | *pch-2(tm1458) II; unc-119(ed3) III; ItIs37 [pAA64; Ppie-1::mCherry::his-58; unc-119 (+)] IV; ItIs38 [pAA1; Ppie-1::GFP::PH(PLC1delta1); unc-119(+)]* |
| BHL596 | *unc-119(ed3) III; ItIs37 [pAA64; Ppie-1::mCherry::his-58; unc-119 (+)] IV; ItIs38 [pAA1; Ppie-1::GFP::PH(PLC1delta1); unc-119(+)]; mdf-1(av19) V* |
| BHL600 | *unc-119(ed3) III; ItIs37 [pAA64; Ppie-1::mCherry::his-58; unc-119 (+)] IV; ItIs52 [pOD379; Ppie-1::GFP::MDF-2; unc-119 (+)]* |
| BHL604 | *pch-2(tm1458) II; unc-119(ed3) III; ItIs37 [pAA64; Ppie-1::mCherry::his-58; unc-119 (+)] IV; ItIs52 [pOD379; Ppie-1::GFP::MDF-2; unc-119 (+)]* |
| BHL608 | *cmt-1(ok2879) I; unc-119(ed3) III; ItIs37 [pAA64; Ppie-1::mCherry::his-58; unc-119 (+)] IV; ItIs38 [pAA1; Ppie-1::GFP::PH(PLC1delta1); unc-119(+)]* |
| BHL664 | *Ppch-2::pch-2::GFP-3XFLAG (blt4 [pCN94]) II; unc-119(ed3) III; ItIs37 [pAA64; Ppie-1::mCherry::his-58; unc-119 (+)] IV* |
| BHL666 | *unc-119(ed3) III; ItIs37 [pAA64; Ppie-1::mCherry::his-58; unc-119 (+)]; mdf-2(tm2190) IV/nT1[qIs51]( IV;V) ItIs38 [pAA1; Ppie-1::GFP::PH(PLC1delta1); unc-119(+)]* |
| BHL883 | *san-1/mdf-3 (ok1580) I; pch-2(tm1458) II; unc-119(ed3) III; ItIs37 [pAA64; Ppie-1::mCherry::his-58; unc-119 (+)] IV; ItIs38 [pAA1; Ppie-1::GFP::PH(PLC1delta1); unc-119(+)]* |
| BHL887 | *cmt-1(ok2879) I; pch-2::GFP-3XFLAG (blt4 [pCN94]) II; unc-119(ed3) III; ItIs44[pAA173; pPie-1::mCherry::PH(PLC1delta1); unc-119(+)]; ItIs37 [pAA64; Ppie-1::mCherry::his-58; unc-119 (+)] IV* |
| BHL888 | *unc-119(ed3) III; ItIs44 [pAA173; pPie-1::mCherry::PH(PLC1delta1); unc-119(+)]; ItIs52 [pOD379; Ppie-1::GFP::mdf-2; unc-119 (+)]; ItIs37 [pAA64; Ppie-1::mCherry::his-58; unc-119 (+)] IV* |
| BHL889 | *pch-2(tm1458) II; unc-119(ed3) III; ItIs44 [pAA173; pPie-1::mCherry::PH(PLC1delta1); unc-119(+)]; ItIs52 [pOD379; Ppie-1::GFP::mdf-2; unc-119 (+)]; ItIs37 [pAA64; Ppie-1::mCherry::his-58; unc-119 (+)] IV* |
| BHL891 | *unc-119(ed3) III; ItIs37 [pAA64; Ppie-1::mCherry::his-58; unc-119 (+)] IV; ItIs24[pAZ132; pPie-1::GFP::tba-2; unc-119(+)]* |
| BHL892 | *pch-2 (tm1458) II; unc-119(ed3) III; ItIs37 [pAA64; Ppie-1::mCherry::his-58; unc-119 (+)] IV; ItIs24 [pAZ132; pPie-1::GFP::tba-2; unc-119(+)]* |
| BHL893 | *san-1/mdf-3 (ok1580) I; unc-119(ed3) III; ItIs37 [pAA64; Ppie-1::mCherry::his-58; unc-119 (+)] IV; ItIs38 [pAA1; Ppie-1::GFP::PH(PLC1delta1); unc-119(+)]* |
| BHL904 | *Ppch-2::pch-2::GFP-3XFLAG (blt4 [pCN94]) II; unc-119(ed3) III; ItIs44[pAA173; pPie-1::mCherry::PH(PLC1delta1); unc-119(+)]; ItIs37 [pAA64; Ppie-1::mCherry::his-58; unc-119 (+)] IV* |
| BHL950 | *pch-2(tm1458) II; unc-119(ed3) III; ItIs37 [pAA64; Ppie-1::mCherry::his-58; unc-119 (+)]; mdf-2(tm2190) IV/nT1[qIs51]( IV;V); ItIs38 [pAA1; Ppie-1::GFP::PH(PLC1delta1); unc-119(+)]* |
| BHL958 | *unc-119(ed3) III; ltIs37 [pAA64; Ppie-1::mCherry::his-58; unc-119 (+)] IV/nT1[qIs51]( IV;V); ltIs38 [pAA1; Ppie-1::GFP::PH(PLC1delta1); unc-119(+)]* |
| BHL964 | *zyg-1(or297) unc-4(e120) II unc-119(ed3) III; ltIs37 [pAA64; Ppie-1::mCherry::his-58; unc-119 (+)] IV; ltIs24[pAZ132; pPie-1::GFP::tba-2; unc-119(+)]* |
| BHL965 | *zyg-1(or297) pch-2(tm1458) unc-4(e120) II; unc-119(ed3) III; ltIs37 [pAA64; Ppie-1::mCherry::his-58; unc-119 (+)] IV; ltIs24[pAZ132; pPie-1::GFP::tba-2; unc-119(+)]* |
| GAL5 | *san-1/mdf-3 (mat5) I; unc-119(ed3) III; ruls57[Ppie-1::GFP::tubulin + unc-119(+)]; ltIs37[pie-1p::mCherry::his-58 (pAA64) + unc-119(+)] IV* |

